# Time Dependent Stochastic mRNA and Protein Synthesis in Piecewise-deterministic Models of Gene Networks

**DOI:** 10.1101/278226

**Authors:** Guilherme C.P. Innocentini, Arran Hodgkinson, Ovidiu Radulescu

## Abstract

We discuss piecewise-deterministic approximations of gene networks dynamics. These approximations capture in a simple way the stochasticity of gene expression and the propagation of expression noise in networks and circuits. By using partial omega expansions, piecewise deterministic approximations can be formally derived from the more commonly used Markov pure jump processes (chemical master equation). We are interested in time dependent multivariate distributions that describe the stochastic dynamics of the gene networks. This problem is difficult even in the simplified framework of piecewise-determinisitic processes. We consider three methods to compute these distributions: the direct Monte-Carlo, the numerical integration of the Liouville-master equation and the push-forward method. This approach is applied to multivariate fluctuations of gene expression, generated by gene circuits. We find that stochastic fluctuations of the proteome and much less those of the transcriptome can discriminate between various circuit topologies.

## 1 Introduction

One of the greatest problems of molecular biology is how single undifferentiated cells give rise to many different cells types, all being genetically identical yet performing different functions. Since the pioneering work of Jacob and Monod [24] it is known that this multiplicity of behaviours is possible because the protein production depends not only on the existence of a gene but also on the quantities of regulatory molecules that can change with the cell type and environmental conditions. The protein production takes place in two steps [2]. First, during a process called transcription, the genetic information from DNA is copied on the messenger RNA (mRNA). Then, during a process called translation, the mRNA is used as template for protein production by the ribosomes. The amount of transcribed mRNA and translated protein, namely the gene expression, can vary from one cell to another for various reasons. One of these reasons is that the gene expression is not a property of a single gene but is a property of a set of interacting genes; a gene network [26]. As per usual in complex systems, the whole is different from the sum of its parts. In this case, a gene network can have many different stable expression levels that correspond to different network attractors [50]. Another reason for expression variability, is the fact that transcription and translation are stochastic [32, 11, 49]. A single cell transcription is often intermittent, periods of strong mRNA production being followed by periods of silence when transcription is stopped [49, 40, 6, 48]. The periods of inactivity may correspond to paused RNA polymerase, incompletely formed activation complexes, transient presence of regulatory proteins and complexes on the DNA, or to transitions between open and condensed chromatin [36]. The durations of the periods of activity and inactivity are random. Furthermore, snapshots of the cell population at various times show a heterogeneous picture in which a significant proportion of the cell population express much lower or much higher than the average protein levels [11, 12]. The observed distributions of the gene expression can deviate significantly from a Gaussian by having skewness [12], heavy tails or multimodality [36].

The gene expression stochasticity can have important biological consequences. In infections and tumours, part of pathogen populations (bacteria, viruses, cancer) can escape drug treatment when they adopt behaviours very different from the average; for instance they may stop growth [10, 3], remodel their metabolism [19], or stop absorbing, or expel drugs [18]. Latency phenomena [43] can be responsible for resistance to treatment and relapse of the disease once the treatment is stopped.

Gene expression stochasticity is also important in synthetic biology. Synthetic biology devices are supposed to have a well defined biological function for given combinations of the input conditions. Therefore, in the absence of error-correction, the precision of biological devices depends critically on the amount of stochasticity [41, 35].

For all these reasons, one needs mathematical tools for predicting quantitatively the amplitude and distribution of gene expression fluctuations in gene networks.

Models of stochastic gene networks represent molecular interactions as discrete events (biochemical reactions) separated by random waiting times. This modeling approach, introduced by Delbruck [1] and further developed by Renyi [44] and Bartholomay [4] covers practically all the aspects of gene/gene interactions but is mathematically and computationally challenging. Indeed, the underlying models are continuous time Markov processes with an infinite or extremely large number of states. The corresponding master equation can be solved exactly only in a limited number of cases, corresponding to single genes without or with much simplified feed-back control [37, 20, 21, 42]. The Monte-Carlo simulation algorithms (Gillespie algorithm [15]) can be very inefficient for simulating the full process, as the number of individual reactions that have to be simulated for a significant change of the system’s state can be tremendously large.

Several types of approximation were used in order to simplify stochastic biochemical networks in order to reduce their simulation time and to facilitate their analysis. These simplifications were possible because the stochastic gene networks have heterogeneous variables and multiple time scales [39]. This heterogeneity comes from the fact that some variables *X*_*D*_ such as DNA/regulatory proteins and complexes/polymerase states are discrete and other variables *X*_*C*_ such as protein and mRNA copy numbers are continuous. The species dichotomy leads to a partition of the biochemical reactions. Although literature provides a number of different ways for reaction partitioning, the four set partition *R*_*D*_, *R*_*C*_, *R*_*DC*_, *R*_*CD*_ (Figure 1) seems to us quite natural. A very similar partitioning was used for rigourously justified approximations in [8, 7].

**Figure 1:**
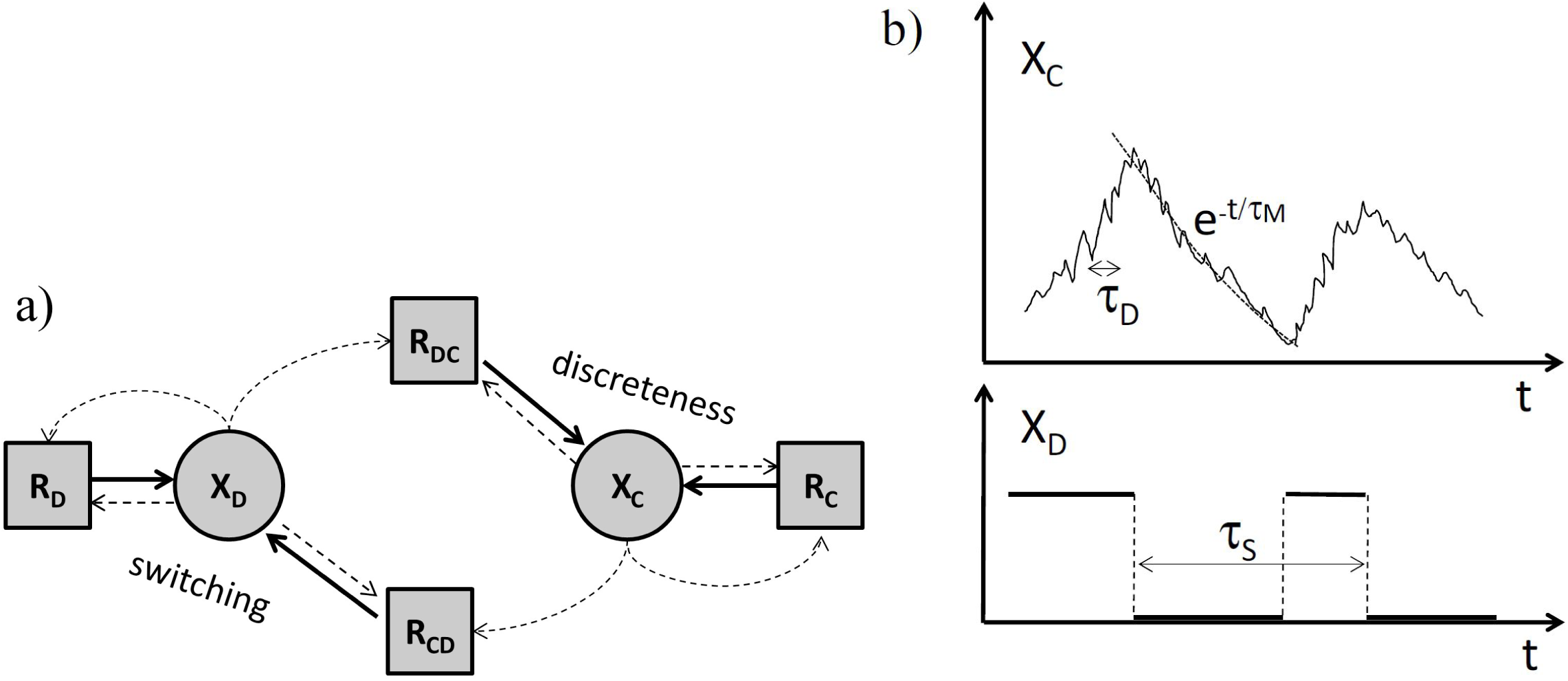
Types of variables and important timescales of stochastic biochemical reactions networks (a) The partition of variables and of the reactions. Dotted line arrows mean that reaction rates depend on the source variables. Continuous line arrow mean that the source reaction acts on the end variable. *R*_*D*_ are reactions acting on discrete variables and whose rates depend on discrete variables, *R*_*C*_ are reactions acting on continuous variables and whose rates depend on continuous variables. *R*_*CD*_, *R*_*DC*_ are reactions acting on discrete and on continuous variables, respectively, and have rates depending on both discrete and continuous variables. (b) Typical trajectories and time scales of continuous and discrete variables.

Two main timescales have to be taken into account in order to find the appropriate approximation. As shown in the Figure 1, the discrete variables switch between a number of discrete states. The characteristic time of this process was called switching time, *τ*_*S*_ [39]. The trajectories of continuous variables are smooth only in the average, but for these variables, the average is a good approximation. The characteristic time of fluctuations of continuous species around their average was named discreteness time, *τ*_*D*_ [39]. The discreteness time scales like 1*/N* where *N* is the copy number of the continuous species.

There are two main classes of approximated models (Figure 2).

**Figure 2:**
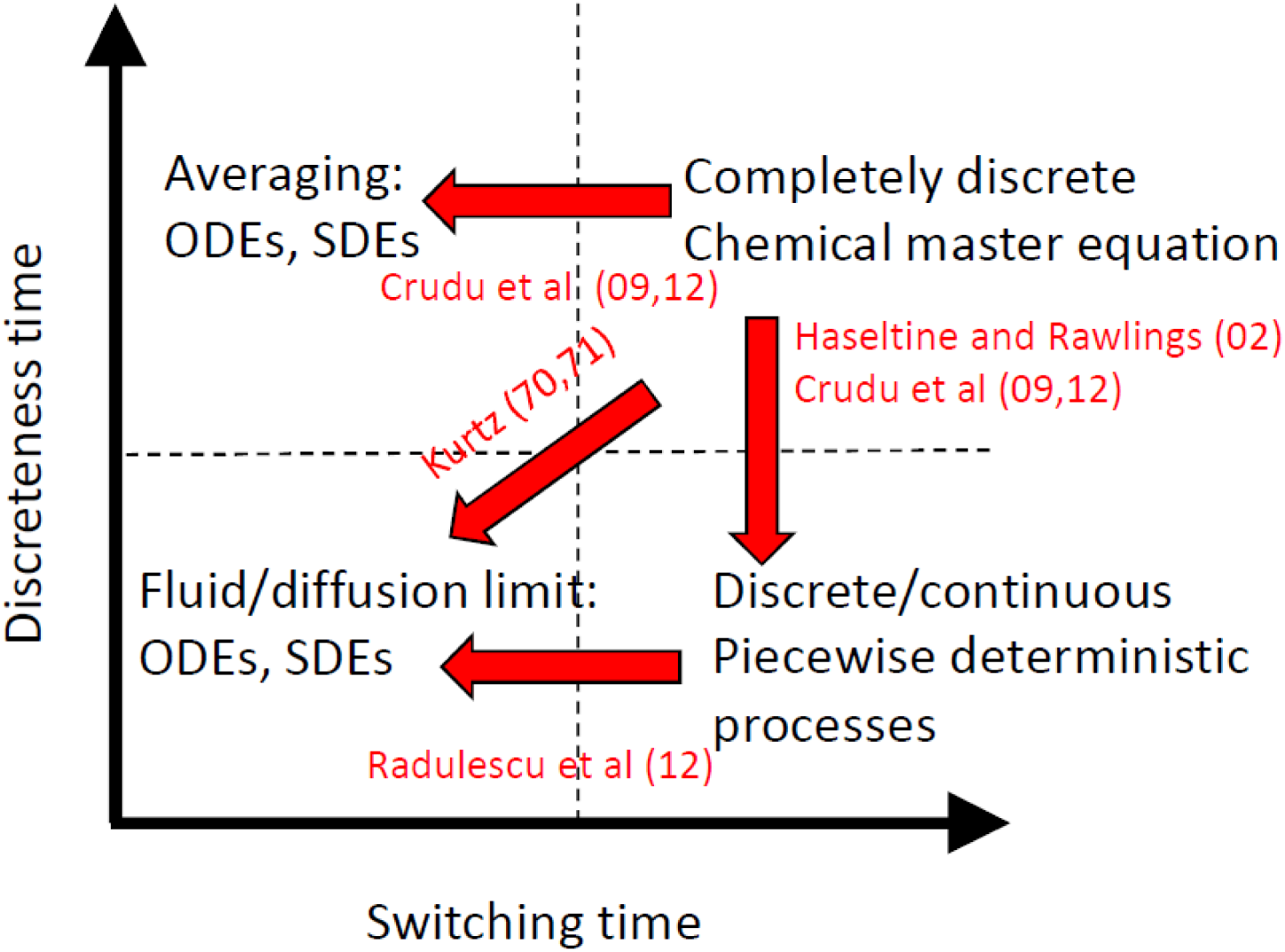
Different approximations of stochastic biochemical reactions networks. The “exact” model is a completely discrete Markov process described by the chemical master equation. Various approximations can be obtained as asymptotic limits when some parameter is very small or very large from the “exact” model or from approximations.

- Deterministic (ordinary differential equations, ODEs) or diffusion (Fokker-Planck) approximations can be applied when the transitions between discrete states are fast and the continuous species have large copy numbers, thus when *τ*_*S*_, *τ*_*D*_ are both small. Then one obtains the deterministic or diffusion limit by applying the law of large numbers or the central limit theorem to Markov processes [28, 29]. This result can be obtained heuristically by the Ω expansion, well known in physics [51]. If the discreteness time is larger than the switching time, similar approximated models can be obtained by using averaging [8, 7]. In both cases the discrete states are homogenized and the approximated model has only continuous variables.
- Piecewise-determinisitic processes (PDP) or hybrid diffusion approximations can be obtained when the discreteness time is small, but *τ*_*S*_ *>> τ*_*D*_. In these approximations there are two types of variables. The dynamics of the continuous variables is described by ODE or stochastic differential equations (SDE), whereas the discrete variables dynamics is described by Markov chains. These approximations were first obtained heuristically by using partial Ω expansions of the master equation [17, 38, 8] and then justified rigorously by using generators and measure theory in [7]. Finally, diffusion approximation was applied to PDP in the limit when the switching time is small, to obtain again deterministic and Fokker-Planck approximations [39].

In this paper we consider the situation when the swiching time is relatively large. In biology, this corresponds to the so-called “random telegraph”[30], “bursting” [40, 12], or “multi-scale bursting” [48] fluctuations. Simply speaking there are some variables (DNA, regulatory complexes and/or RNA polymerase states) that have ON-OFF or multiple states Markov chain dynamics. The discrete variables control the ODE dynamics of the continuous variables (mRNA, proteins). The underlying approximation is piecewise-deterministic, because the ODEs change each time when the discrete variables perform a transition (see Figure 1).

In our PDP models of gene networks each gene promoter is described as a finite state Markov chain. The transition rates between states of the promoter depend on the expression levels of the regulating genes. The promoter triggers synthesis of gene products with intensities depending on its state. We represent the state of a gene network as a random vector whose components indicate the promoter state and the copy numbers of all the mRNAs and proteins in the network. The dynamics of the gene network will be represented as the time-dependent multivariate distribution of this random vector. These distributions contain information about the randomness of each gene but also about correlation among genes and promoter states. We present three methods to compute time dependent multivariate distributions for such models: the direct simulation of the PDP process (the Monte-Carlo method), the numerical solutions of the Liouville-master equation, and the push-forward method. The first method is stochastic, whereas the last two are entirely deterministic (don’t use random number generators). The PDP Monte-Carlo method combines the numerical simulation of the Markov chain of promoter states with symbolic solutions of the deterministic ODEs and is much faster than the direct Gillespie method [15].

This paper is structured as follows: In Section 2 we recall the justification of piecewise-deterministic approximation using the partial Ω expansion. In Section 3 we introduce a class of piecewise-deterministic models covering gene network. In Section 4.1 we discuss the Monte-Carlo methods for piecewise-deterministic models. In Section 4.2 we discuss the Liouville-master equation approach. In Section 4.3 we discuss the push-forward method. In Section 5 we briefly discuss possible applications of these methods to extracting information from mRNAs’ and proteins’ spectra of fluctuations.

## 2 Partial omega-expansion and the Liouville-master equation

The dynamics of stochastic biochemical networks can be described by a pure jump Markov process [1, 44, 4] (for a general presentation of Markov jump processes see [14]). The state of the network is a vector *X ∈* ℕ^*n*^ whose components represent copy numbers of molecules of various species. Each biochemical reaction is defined by a stoichiometric (jump vector) *γ*_*i*_ *∈ ℤ*^*n*^, *i ∈ ℛ* where *ℛ* is the set of biochemical reactions in the model. The occurrence of the reaction *i* is represented as a jump in the system’s state *X → X* + *γ*_*i*_. Finite, discrete states of gene promoters can be represented by extending the meaning of species to include “places” with finite values of the copy numbers. For instance, on/off gene promoters can be represented by using two places *P*_on_ and *P*_off_ with two possible occupancies 0 or 1. The transitions from *OFF* to *ON* and back can be represented by a reversible reaction that consumes a particle on the place *P*_off_ and produces a particle on the place *P*_on_.

In the Markov pure jump representation the dynamics of the system is a series of jumps separated by exponentially distributed waiting times [14, 38]. The number of reactions of the type *i* occurring in the average per unit time is given by the propensity (or rate) function *X → V*_*i*_(*X*; *µ*), where *µ* are kinetic parameters. The time between successive reactions is exponentially distributed with an average (Σ _*i∈ ℛ*_ *V*_*i*_)^−1^ and the next reaction is *i* with probability *V*_*i*_*/*(Σ_*i∈ℛ*_*V*_*i*_). We are interested in the multivariate distribution *p*(*X, t*) representing the probability to be in state *X* at time *t*. This satisfies the time dependent master equation:

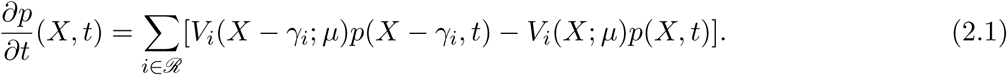

The van Kampen Ω (system size) expansion [51] or equivalently, the central limit theorem [28, 16] lead, in the first order, to deterministic (ODE) approximations and to diffusion (Fokker Planck) approximations, in the second order. The partial omega expansion consists in applying the Ω expansion only to species that are produced in sufficiently large copy numbers [7, 8]. The large copy numbers species denoted *X*_*C*_ are called continuous, because the biochemical reactions change their values gradually, by the accumulation of a large number of small steps. Other species are present only in a few copies per cell (these include the “places” describing promoter states). We denote these species *X*_*D*_ and call them discrete. This leads to a decomposition of the state vector as *X* = (*X*_*D*_, *X*_*C*_) and of all stoichiometric vectors as 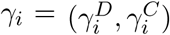, corresponding to discrete and continuous species coordinates. The interactions among discrete and continuous species are suitably described by a partition of the reactions in four sets *ℛ* = *ℛ*_*D*_ *∪ℛ*_*DC*_ *∪ ℛ*_*CD*_ *∪ ℛ*_*C*_ [7, 8, 39].

The reactions *ℛ*_*D*_ act on *X*_*D*_ (the corresponding *γ*_*i*_ have non-zero coordinates on discrete species, 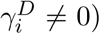 and have propensities depending on *X*_*D*_ only. The reactions *ℛ*_*C*_ act on *X*_*C*_ (the corresponding *γ*_*i*_ have non-zero coordinates on continuous species, 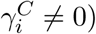 and have propensities depending on *X*_*C*_ only. The reactions *ℛ*_*DC*_, *ℛ*_*CD*_ act on *X*_*C*_ and *X*_*D*_, respectively, and their propensities depend on both *X*_*D*_ and *X*_*C*_.

In this paper we consider gene network models. For each gene, we model the transitions between promoter states, as well as other processes such as transcription, translation, protein folding, and protein and mRNA degradation. We will consider that the mRNA molecules and proteins are in sufficiently large copy numbers to justify continuous approximations. The only discrete variables are in this case the promoter states. The set *ℛ*_*D*_ contains transitions between discrete promoter states whose rates do not depend on regulatory proteins. The set *ℛ*_*CD*_ contains transitions between promoter states whose rates depend on concentrations of regulatory proteins. The set *ℛ*_*C*_ contains translation, maturation (folding), degradation reactions. The set *ℛ*_*DC*_ contains transcription initiation reactions that depend on the promoter state.

We further consider that the copy numbers of continuous species *X*_*C*_ and the propensities of reactions in the sets *ℛ*_*C*_ and *ℛ*_*DC*_ are “extensive”, in other words, scale with the system size Ω, *X*_*C*_ = Ω*x*_*c*_, *V*_*i*_ = Ω*v*_*i*_(*x*_*c*_, *X*_*D*_), for *i ∈ ℛ*_*DC*_ *∪ℛ*_*C*_, whereas the propensities of reactions in *ℛ*_*D*_ and *ℛ*_*CD*_ are considered “intensive” and do not scale with Ω. The Ω dependence is not only a useful mathematical tool, but has also a biological meaning. In proliferating cells, the protein concentrations are important for biochemical reactions and should be maintained by synthesis reactions. This is only possible if the propensities of synthesis reactions (including mRNA synthesis) scale with the size. Rates of monomolecular reactions consuming continuous reactants (for instance, degradation reactions) are proportional to the reactant copy number that scale with the size. Rates of switching reactions between discrete promoter states do no scale with size, unless they are proportional to copy numbers of activator or repressor proteins. For a more complete discussion of these scaling relations we refer to [8].

In rescaled variables, the master equation reads

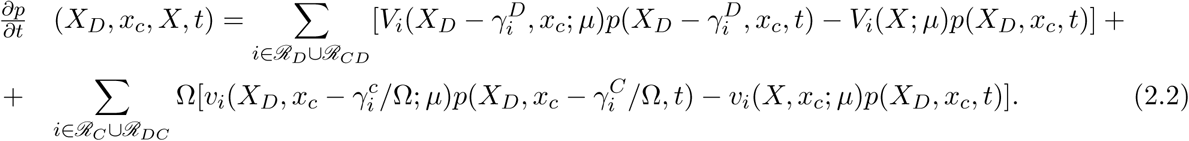

Using the first order Taylor series expansion 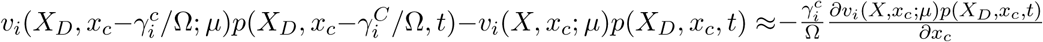 we obtain the Liouville-master equation

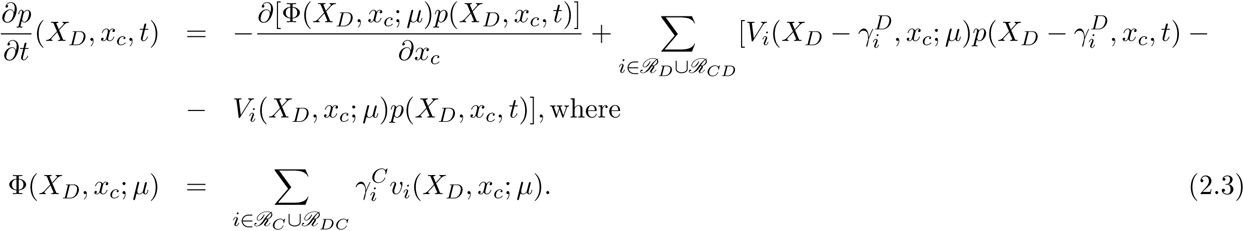

The Liouville-master equation has been used in various fields; in statistical physics of quantum systems [27, 13, 5] or with a different name in statistics and operations research [9]. It describes the time-dependent distribution of a piecewise-deterministic model. Indeed, conditionally on *X*_*D*_ (considering that the discrete variables are fixed) the continuous variables *x*_*c*_ satisfy deterministic, ODEs:

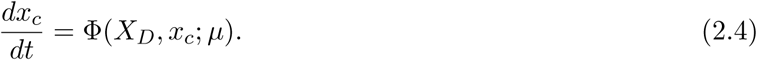

The discrete variables follow a pure jump Markov dynamics defined by the stoichiometric vectors 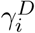 and the propensities *V*_*i*_(*X*_*D*_, *x*_*c*_; *µ*), *i ∈ ℛ*_*D*_ *∪ ℛ*_*CD*_. The noise in the system is produced by the stochastic transitions of the discrete variables. The probability distribution of the continuous variables is advected (transported by the flow defined by the ODEs, see Figure 3) by the deterministic flow Φ(*X*_*D*_, *x*_*c*_; *µ*) in the continuous variables space; this explains the advection term 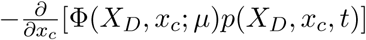 in the Liouville-master equation.

**Figure 3:**
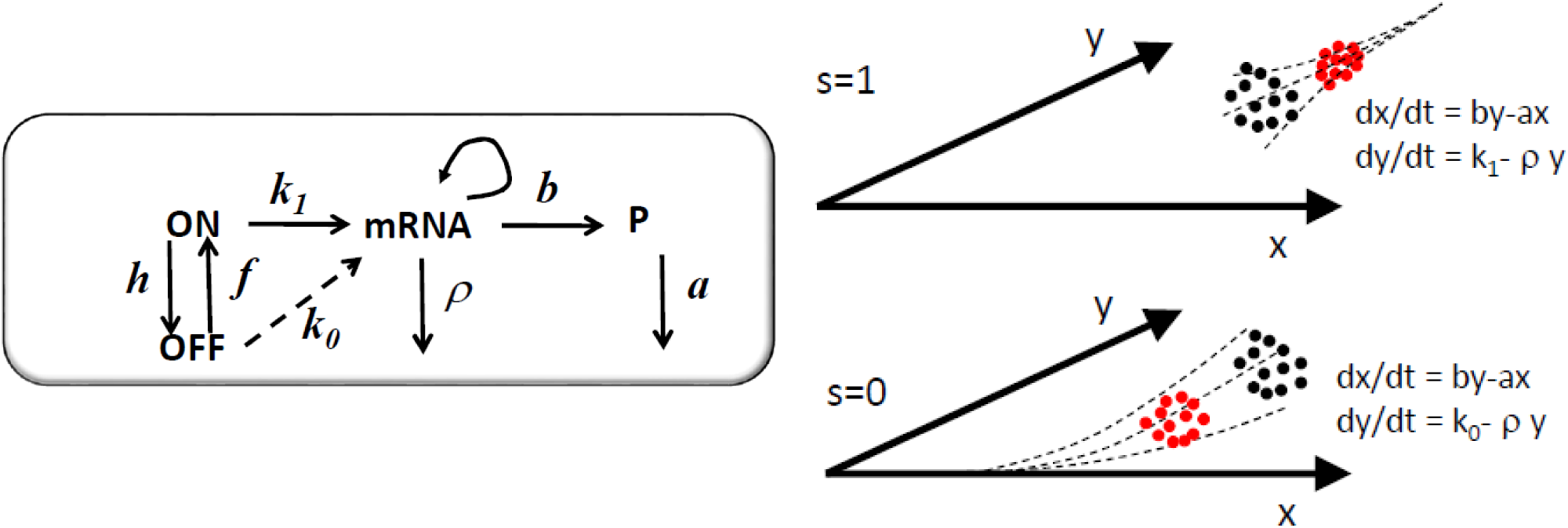
Model of a single gene controlled by a two state (ON/OFF) promoter. In the graphical representation of the model, the arrow from mRNA to mRNA indicates that mRNA is recovered after the transcription reaction. Between two successive jumps of the discrete variables, a probability distribution of the continuous variables x (protein) and y (mRNA) represented as black dots is continuously advected by one of the two possible deterministic flows to a distribution represented by red dots. The *s* = 1 flow pushes the system towards the high expression attractor *x* = (*bk*_1_)*/*(*aρ*), *y* = *k*_1_*ρ* and the *s* = 0 flow pushes the system towards the low expression attractor *x* = (*bk*_0_)*/*(*aρ*), *y* = *k*_0_*ρ*, where *k*_0_ *<< k*_1_. The systems chooses alternately and stochastically between these two flows.

## 3 PDP models of gene circuits

In this section we introduce a family of PDP models that can be used to represent gene networks. They are a special, simplified case of the class of models defined by (2.3). The main simplification is that the propensities of reactions in *ℛ*_*DC*_ depend on *X*_*D*_ and do not depend on *x*_*C*_.

As an example, we consider a gene circuit with dichotomous noise. This model is made of *n*_*g*_ genes, each one controlled by a promoter with 2 states; ON and OFF. The continuous variables are *x*_*i*_ and *y*_*i*_; the protein and mRNA level for each gene *i*, respectively. The discrete variables are two values variables *s*_*i*_ *∈ {*0, 1*}* representing the promoter states (0 stands for OFF and 1 stands for ON). In this model, the set *ℛ*_*DC*_ consists of transcription initiation reactions. Their rates depend on the promoters states (ON of OFF) but do not depend on the protein or mRNA levels. The set *ℛ*_*CD*_ *∪ ℛ*_*D*_ consists of reversible transitions between the states ON and OFF. The corresponding transition rates are constant when the gene *i* is constitutive, namely these rates are *f*_*i*_ from ON to OFF and *h*_*i*_ from OFF to ON. When the gene *i* is activated by the gene *j* the transition rate from OFF to ON is *f*_*i*_*x*_*j*_, whereas when the gene *i* is repressed by the gene *j* the transition rate from ON to OFF is *h*_*i*_*x*_*j*_. The discrete variables’ dynamics is thus a Markov chain with the state set *M* = *{*0, 1*}*^*ng*^. It is convenient to relabel the states from 1 to 2^*ng*^ using the lexicographic order on *M*. For instance, two gene circuits have, in order, the states 1: (0, 0), 2: (0, 1), 3: (1, 0), 4: (1, 1). Also, instead of using reaction propensities, we equivalently provide a transition rate matrix *S* whose elements are the transition probabilities per time unit between states. For instance, for a two gene circuit where the first gene is constitutive and the second gene is activated by the first one, we have:

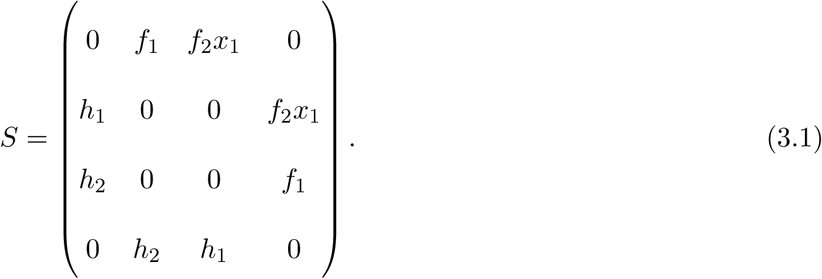

The mRNA and protein variables follow ODE dynamics

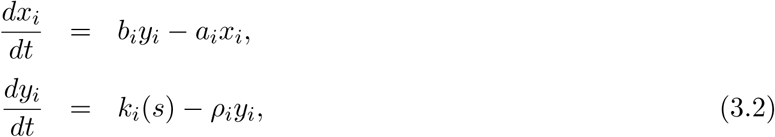

where *ρ*_*i*_, *b*_*i*_, *a*_*i*_, *i ∈* [1, *n*_*g*_] represent mRNA degradation, translation, and protein degradation rates for the gene *i*, respectively, and *s* = (*s*_1_, *s*_2_, *…, s*_*n*_*g*) is the state of the Markov chain, such that

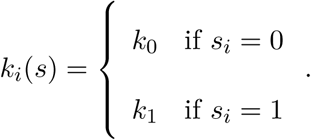

For the sake of illustration let us consider the simple model of a single constitutive gene controlled by a two state (ON/OFF) promoter. We denote the states of the promoter by 1 and 0 respectively. The transition rate from 0 to 1 is *f* and from 1 to 0 is *h*. The protein and mRNA concentrations are *x* and *y*, respectively. The transcription initiation rate in the state 1 is *k*_1_ and in the state 0 is *k*_0_ *<< k*_1_. The translation rate is *b*. The mRNA and protein degradation rates are *ρ* and *a*, respectively. The master-Liouville equation reads

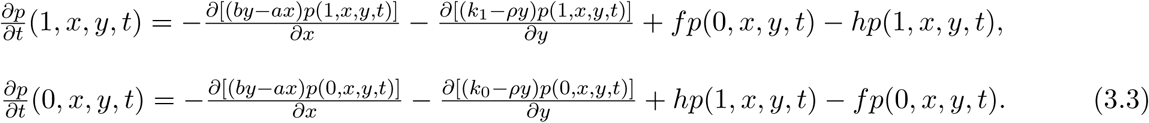

The protein and mRNA concentrations follow the ODEs

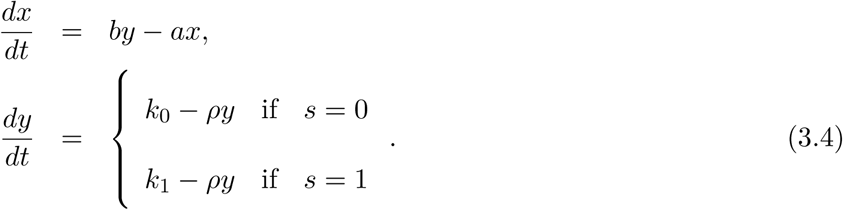

The probability distribution of the promoter state *s* results from the dynamics of the two state Markov chain

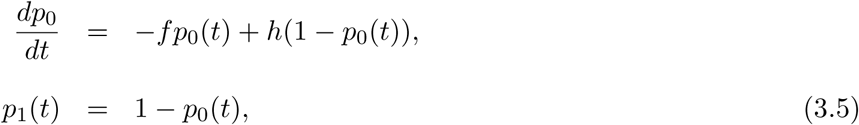

where *p*_0_ = ℙ [*s* = 0] = ∫ *p*(0, *x, y*) *dxdy*, *p*_1_ =ℙ [*s* = 1] = ∫ *p*(1, *x, y*) *dxdy*.

We also define the asymptotic occupancy probabilities 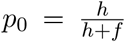 and 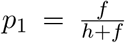, representing the probabilities, at steady state, that the promoter state is OFF and ON, respectively.

The single constitutive gene model and the advection fluxes of the Liouville-master equation are illustrated in Figure 3. More complex, two gene circuits models are represented in Figure 4 and their Liouville-master equations are given in the Appendix 1.

**Figure 4:**
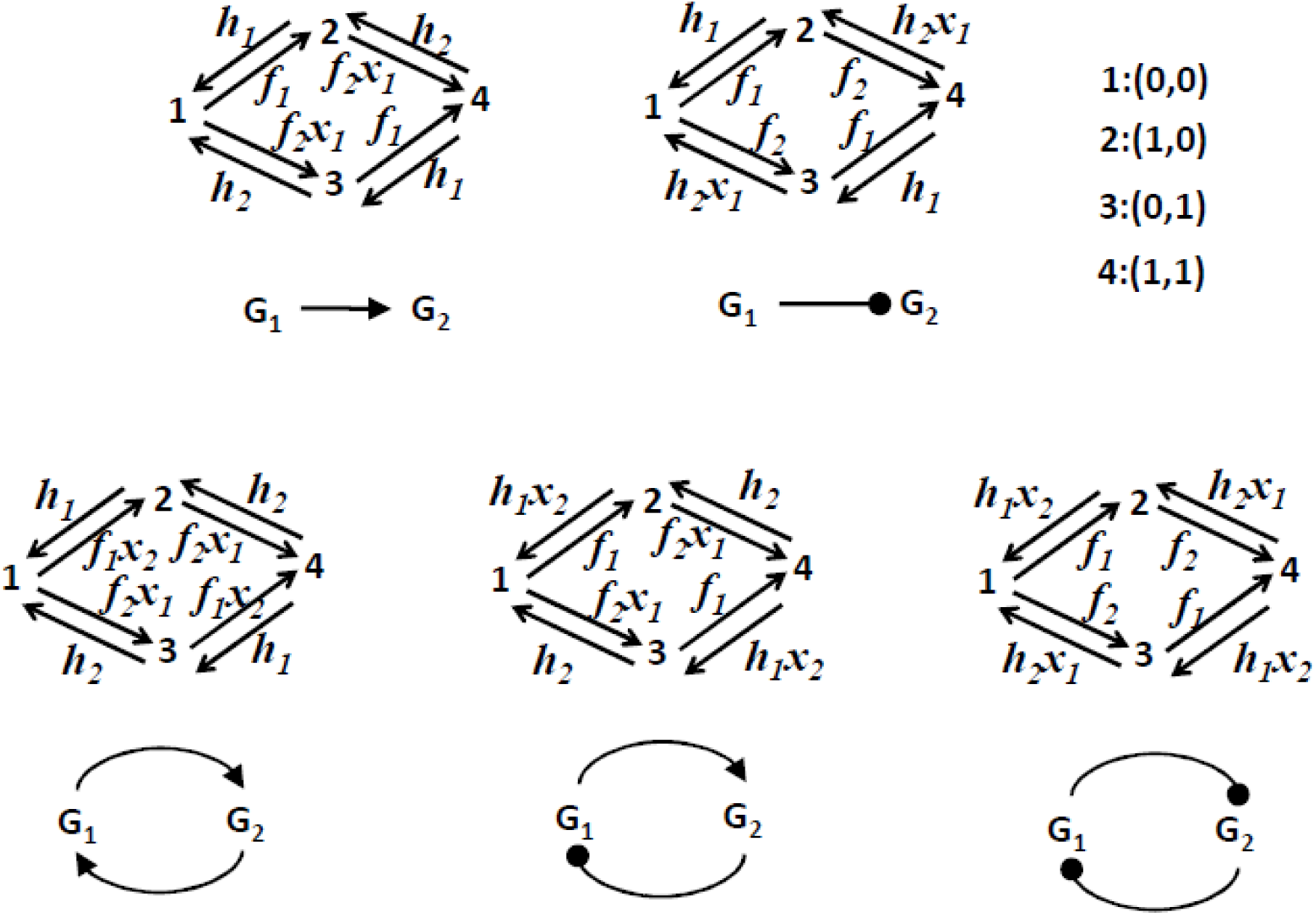
Two gene circuits models. Only the discrete (Markov chain) part of the dynamics is represented. These models have 4 discrete states as each one of the two promoters can be ON or OFF. The transition rates between states are either constant or functions of the levels of proteins *x*_1_ and *x*_2_. We consider that transcription, translation and degradation parameters *k*_0_, *k*_1_, *ρ*, *a*, *b* are the same for the two genes.

## 4 The numeric methods for solving the Liouville-master equation

In this section we compare several numerical methods for solving the Liouville-master equation in the context of gene networks models. In order to quantify the relative difference between methods we use the *L*^1^ distance between distributions. More precisely, if *p*(*x*) and 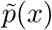 are probability density functions to be compared, the distance between distributions is

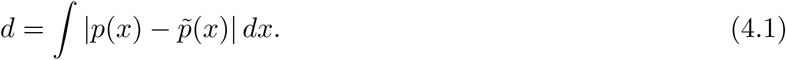

### 4.1 The PDP Monte-Carlo method

The PDP Monte-Carlo method is based on the direct simulation of the PDP process. A simple algorithm has been proposed in [8]. For the sake of completeness we recall here the main steps of this algorithm.

(1) Set the initial state condition *s* = *s*_0_, *x* = *x*_0_, *y* = *y*_0_, *t* = 0.

(2) Generate a random variable *u* uniformly distributed in [0, 1],

(3) Integrate the system of differential equations obtained by adding to (3.2) the equation for the survival function *F* of the waiting time to the next Markov chain transition

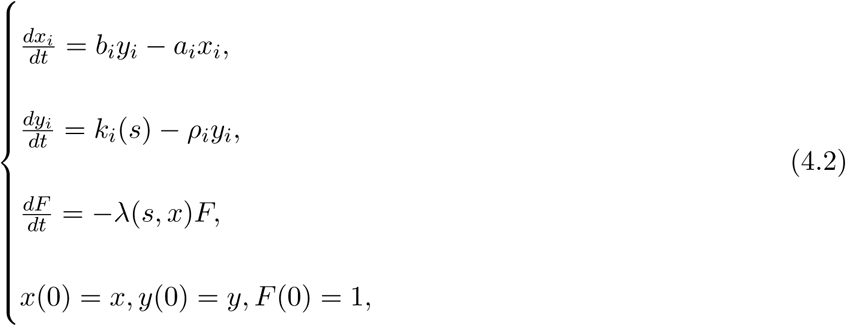

between 0 and *τ* with the stopping condition

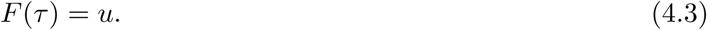

Here *λ*(*s, x*) is the sum of transition probabilities in the row corresponding to the state *s* of the transition rates matrix *S*.

(4) Generate a second uniform variable *v* and use it to find the next Markov chain state *s*_*next*_. The decision for the next discrete state is made in the same way as in the Gillespie algorithm [15].

(5) Change the system state *s* = *s*_*next*_, *x* = *x*(τ), y = y(τ), and the time t = t + τ.

(6) Reiterate the system from (2) with the new state until a time *t*_*max*_ previously defined is reached.

(7) Store samples of x and y at various predefined times.

(8) Reiterate the system from (1) a large number of times. Perform distribution estimates using the resulting statistical ensemble.

A major improvement with respect to [8] can be obtained by using symbolic solutions of (4.2) (see the Appendix 2 for the symbolic solutions) and solving numerically the nonlinear stopping condition (4.3).

### 4.2 The finite difference (FD) Liouville-master method

The Liouville-master equation is a system of linear, partial differential equations (PDEs) for the probability distributions of the mRNAs and proteins for various states of the gene promoters. The number of PDEs is equal to the number of distinct discrete states of the gene promoters, i.e. it is 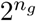, where *n*_*g*_ is the number of genes in the network.

These equations have to be integrated with boundary conditions in order to control possible mass accumulation on the integration domain boundaries. The boundary conditions are obtained by setting to zero the advection fluxes pointing towards the boundary. For the one-gene model (3.3), the integration domain is 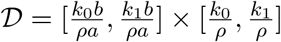 and the boundary conditions read

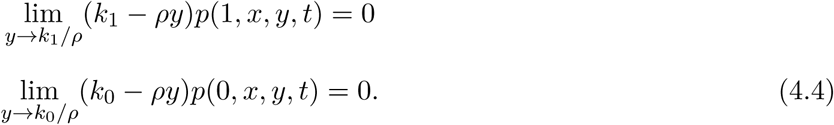

One can note the limit version of the boundary conditions allow divergence of *p*_0_, *p*_1_ on the boundaries *y* = *k*_0_*/ρ* or *y* = *k*_1_*/ρ*, respectively. For boundaries different than *y* = *k*_0_*/ρ* or *y* = *k*_1_*/ρ*, advection fluxes point towards the interior of the integration domain and one can simply impose zero boundary conditions *p*_0_ = *p*_1_ = 0.

As noticed by Marc Kac in a very instructive paper about a piecewise-deterministic random walk [25], in contrast to the Fokker-Planck equations that describe “normal” random-walk diffusion and are parabolic, the Liouville-master equation has hyperbolic nature. General properties of hyperbolic equations, such as finite propagation velocity of perturbations and existence of sharp fronts apply to our equations as well. For our problem, the front discontinuities occur at the domain boundaries and they are handled by the boundary conditions (4.4). Hyperbolicity properties are mainly visible at slow switching and should disappear at fast switching when the Liouville-master equation can be well approximated by a Fokker-Planck equation [39].

In this paper we have used a finite-difference predictor-corrector scheme [47] to compute the solution of the Liouville-master equation. In Figure 5 we compare the distributions for mRNA and proteins resulting from the constitutive gene model (3.3) with the Monte-Carlo simulation of the model. The comparison is quantitative and uses the distance defined by (4.1). In all our computations, the asymptotic occupancy probability is one half. For slow switching, the mRNA distribution is bimodal, with discontinuities at the modal values *k*_0_*/ρ* and *k*_1_*/ρ* values, where the probability density diverges on one side and is zero on the other side. The bimodality resulting from slow switching is well understood and signalled in many other places in the literature (see for instance [45, 23]). We can emphasize that the discontinuity of the solution is a consequence of the hyperbolicity of advection fluxes. A parabolic diffusion flux would not be able to build up such discontinuities and this can be seen in the fast switching distributions that are continuous and unimodal. Interestingly, the protein distributions are unimodal in both cases: slow and fast. A unique discontinuity can be observed at short times in the protein distribution, for a slowly switching promoter.

**Figure 5:**
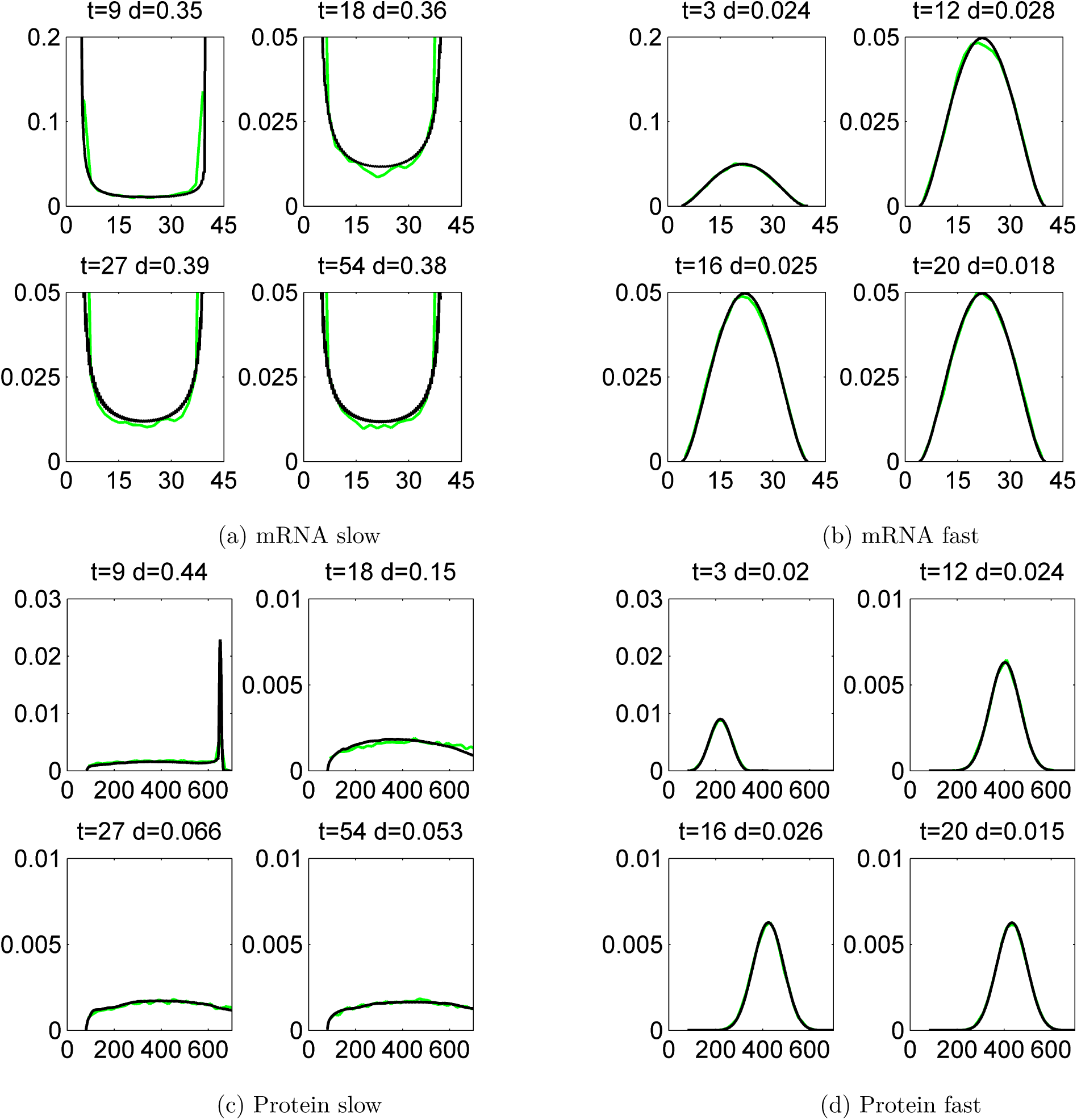
Histograms of protein and mRNA copy numbers for a single gene, produced by the Monte-Carlo method (green lines) and by the finite difference Liouville-master equation method (black lines). The initial data has half-normal bivariate distribution whose mode is *x*_1_ = *x*_2_ = *y*_1_ = *y*_2_ = 0. The gene parameters are *ρ* = 1, *k*_0_ = 4, *k*_1_ = 40, *a* = 1*/*5, *b* = 4, *p*_*a*_ = 0.5, and *ε* = 0.5, *ε* = 5.5, for a slow, and a fast switching gene, respectively. The comparison is quantified by the distance *d* defined by (4.1).

The bivariate mRNA vs. protein distributions are shown at steady state in Figure 6. The mRNA is positively correlated to the protein as it should be. Interestingly, for slow switching, cells close to half protein maximum expression have strongly bimodal mRNA expression, either vanishing or maximum. This rather unintuitive behaviour results from the difference in lifetime between the mRNA and protein molecules. The mRNA molecule has a short life and can, for slow switching, oscillate between very low and maximum values. The protein is much more stable and integrates these oscillations over a large lifetime. This explains why a cell with half protein maximum copy number can have extremely variable mRNA copy numbers.

**Figure 6:**
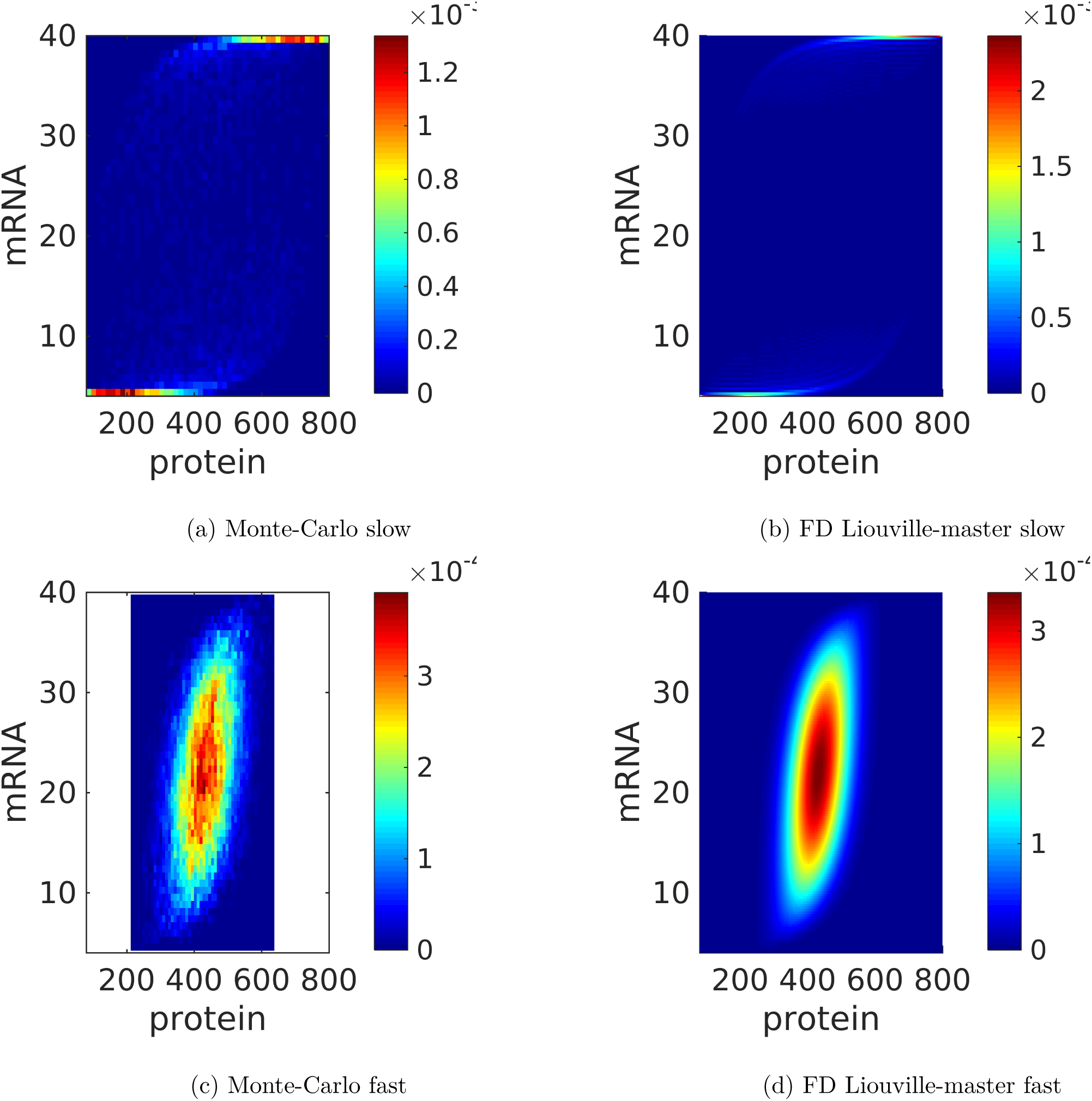
Steady state distributions of protein and mRNA copy numbers for a single gene, produced by the Monte-Carlo method and by the finite difference (FD) Liouville-master equation method. The parameters are those used in Figure 5. The probability to color map relation is linear.

### 4.3 The push-forward method

To introduce the method let us consider the example of a two gene circuit in which the first gene is constitutive and activates the second gene. In other words, the first gene is a transcription factor of the second one.

The push-forward method for gene networks was first introduced in [22]. This method combines Master Equations (MEs) and Random Differential Equations (RDEs). The ME provides the time evolution of the probability distribution of discrete variables. With respect to [22] where the discrete variables were the promoter ON/OFF states and the mRNA copy numbers, here we consider that the discrete variables are solely the promoter’s ON/OFF states. The RDEs are differential equations for the mRNAs and proteins in which the promoter states are considered random parameters. The coupling of ME and RDE is another equivalent way to define the piecewise-deterministic process. When the ME is not dependent on the RDE our models are of the same type as those discussed by Mark Kac in relation to the telegrapher’s equation [25].

Let us introduce our model beginning with the MEs describing the dynamics of the first switch,

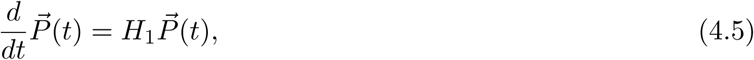

where, 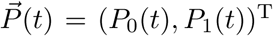 is the probability occupancy vector whose entries are the probabilities to find the first switch in the OFF state (*P*_0_(*t*)) or in the ON state (*P*_1_(*t*)). The infinitesimal stochastic matrix *H*_1_ is given by:

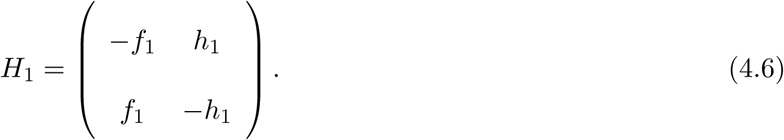

This is a basic telegraph process where the rates *f*_1_ and *h*_1_ are time independent and control the reaction OFF (state 0) to ON (state 1) and ON (state 1) to OFF (state 0), respectively. The equation describing mRNA dynamics is a Random Differential Equation (RDE) given by

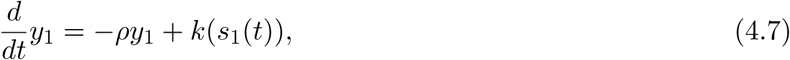

where *y*_1_ is a random variable representing the copy number of mRNA in the cell coming from the first gene. The random variable *s*_1_(*t*) follows the switch statistics, meaning that with probability *P*_1_(*t*), *s*_1_(*t*) = 1 at time *t* and *s*_1_(*t*) = 0 with probability *P*_0_(*t*), again at time *t*. The production rate of mRNA is a function of the random variable *s*_1_(*t*) following

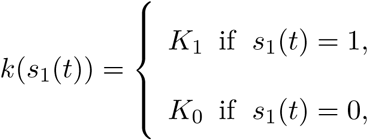

where *K*_1_ is the highest level of mRNA production and *K*_0_ is the basal one. The third equation describing the activity of the first gene is for the random variable representing the protein density associated to it;

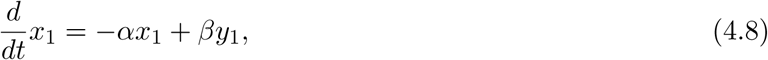

where *α* is protein degradation rate and *β* is the translation rate. The last equation for the coupled gene model is the one governing the probability occupancy of the second gene,

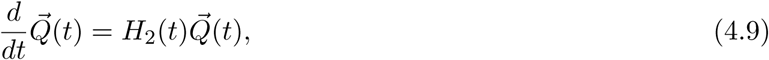

where 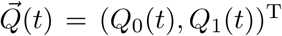 encodes, in its entries, the information about the probability to find the second gene ON (*Q*_1_(*t*)) or OFF (*Q*_0_(*t*)). The matrix *H*_2_(*t*) is given by

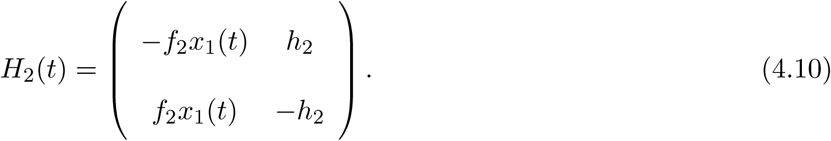

In the model at hand, the main source of stochasticity is the switching ON and OFF of the gene. This noise is transmitted to mRNA synthesis process through the rate *k*(*s*_1_(*t*)) which is a function of a random variable (*s*_1_(*t*)) and, so, a random variable itself. The first step of the push-forward method is to compute the time dependent distribution probability of mRNA molecules *y*_1_(*t*) (which is perturbed by the random variable *s*_1_(*t*)) once the probability distribution of the perturbation is known. To do so, we begin by presenting the solutions of Eqs (4.5),

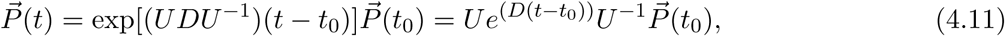

where 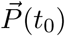 encodes the initial configuration (given at *t* = *t*_0_) of the switch, and the matrices are

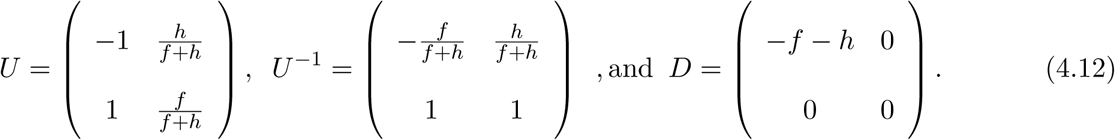

Explicitly, the solutions are given as

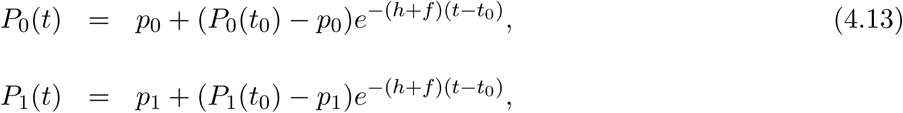

where *p*_0_ = *h/*(*f* + *h*) and *p*_1_ = *f/*(*f* + *h*) are the asymptotic occupancy probabilities to find the gene OFF and ON, respectively. Going on, we present the formal solution of the RDE governing mRNA dynamics,

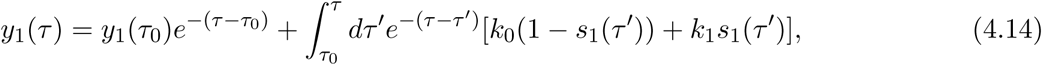

where we have rescaled time *t* by the mRNA degradation rate and introduced the new time parameter, *τ* = *tρ*, and also the dimensionless parameters *k*_0_ = *K*_0_*/ρ* and *k*_1_ = *K*_1_*/ρ*. Note that the integral (4.14) is a basic Riemann integral, such that, if we consider a sufficiently fine partition [*τ*_0_, *τ*_1_, *…, τ*_*N-*1_, *τ*_*N*_] of the integral interval [*τ*_0_, *τ*], where *τ*_*N*_ = *τ*, we can assume that *s*_1_(*τ* ^*l*^) is constant inside each specific partition [*τ*_*j*_, *τ*_*j*+1_] and the integral in (4.14) is approximated by

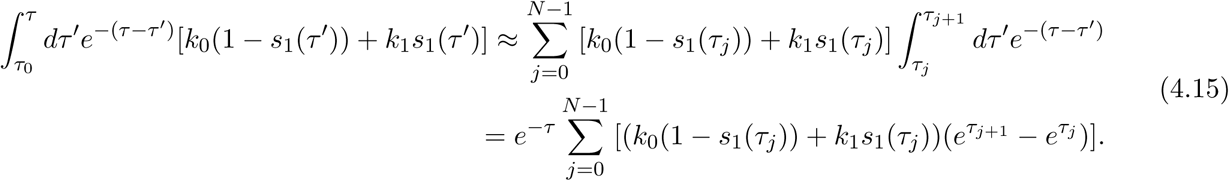

It is worthwhile to remember that *s*_1_(*τ*) is a jumping process assuming, at each instant of time, just one of the values, 0 or 1, with probability *P*_0_(*τ*) or *P*_1_(*τ*), respectively. Now the solution of *y*_1_(*τ*) is given as a function of sequences of *s*_1_(*τ*_*j*_), with *j* = 0, *…, N -* 1. These sequences are strings of zeros and ones and we must consider all of them. For instance, if the number of partitions is *N*, we will have 2^(*N -*1)^ possible sequences and each one will lead to a different value of *y*_1_(*τ*). The remaining task is to assign the correct weight for each one of these values of *y*_1_(*τ*). This is achieved by using the occurrence probability of each possible string of zeros and ones defining the values of *y*_1_(*τ*). The occurrence probability is given by the joint distribution probability of having *s*_1_(*τ*_*j*_) = 0 or 1 at time *τ*_*j*_ for *j* = 0, *…, N -* 1 which is

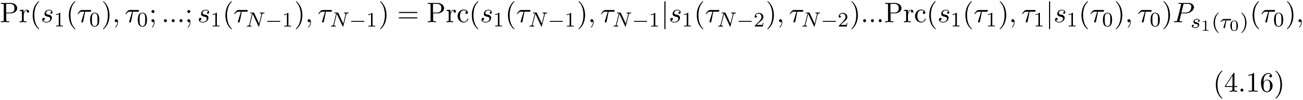

where *P*_*s*_1_(*τ*_0)_(*τ*_0__) is the occupancy probability to find the gene in state *s*_1_(*τ*_0_) at time *τ*_0_, as per (4.13), and Prc(*… | …*) stands for the conditional probability for the telegraph process. For instance,

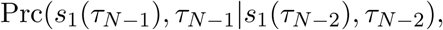

encodes the transition probability from state *s*_1_(*τ*_*N-*2_) to state *s*_1_(*τ*_*N-*1_) during the time interval *τ*_*N-*1_ *- τ*_*N-*2_ knowing that at *τ*_*N-*2_ the system was with probability one in state *s*_1_(*τ*_*N-*2_). The conditional probabilities for the telegraph process, which is a Markovian process, are easily obtained by chosen the appropriate initial configuration:

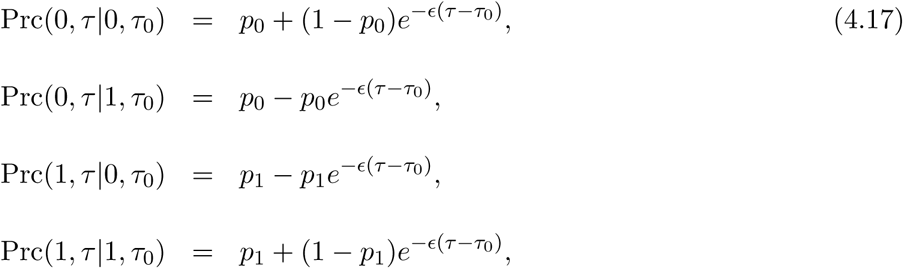

where the new parameter *ε* = (*f* + *h*)*/ρ* measures the flexibility of the switch. With the conditional probabilities and (4.15) at hand, we can calculate the time dependent probability distribution for the mRNA density. We have considered two examples: one is the fast switch regime (*ε >* 1) and, the other, slow switching (*ε <* 1).

Protein distribution in time can be obtained in the same fashion as the one for mRNA. The general solution for protein density is:

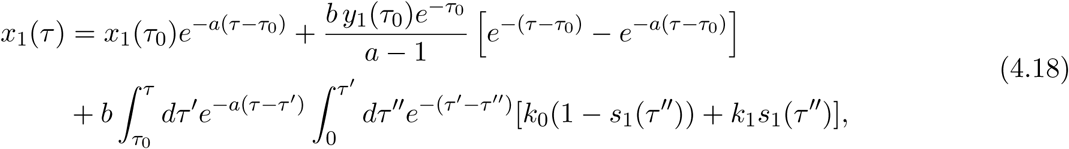

where the rescaled parameters are given by *a* = *α/ρ* and *b* = *β/ρ*. The push-forward method can be applied to obtain the time dependent distribution probability for protein density, in an analogous way as for mRNA. The integral that must be partitioned is that over *τ* ^*l*^, in the interval [*τ*_0_, *τ*]

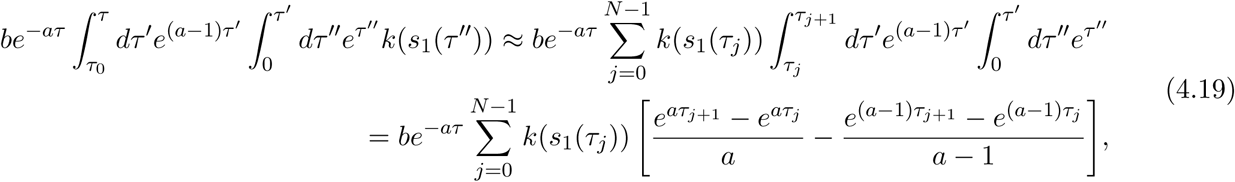

where, we have used the definition *k*(*s*_1_(*τ*)) = *k*_0_(1 *- s*_1_(*τ*)) + *k*_1_*s*_1_(*τ*) to simplify the notation. As before, we have illustrated our method by calculating the protein density for the same two regimes of switch flexibility.

To analyze the influence of the first gene on the second one we have assumed that the action of the first gene is to activate the second (see (4.10)). To do so, instead of solving the RDE describing the activity of the second gene (it is an RDE because the perturbation *x*_1_(*τ*) is a random variable) we have analyzed the mean value of the occupancy probability of the second gene whose dynamics is given by

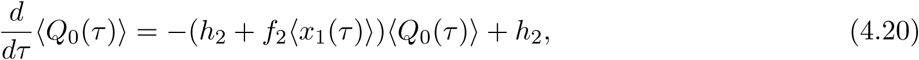

and 〈*Q*_1_(*τ*)〉 = 1 *- 〈Q*_0_(*τ*)〉. The general solution for 〈*Q*_0_(*τ*)〉 is given by

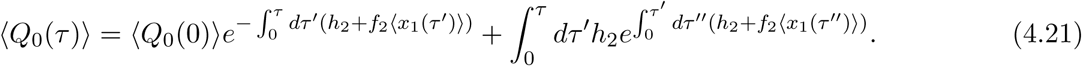

In Appendix 3 we show, in detail, how to obtain the exact functional shape of 〈*x*_1_(*τ*)〉. Nevertheless, its structure is 〈*x*_1_(*τ*)〉 = *r*_0_ + *r*_1_*e*^*-τ*^ + *r*_2_*e*^*-aτ*^ + *r*_3_*e*^*-Eτ*^ and, because of this, the integral in (4.21) cannot be evaluated analytically and a numerical evaluation must be performed. This will also be the case for the conditional probabilities that will be expressed as

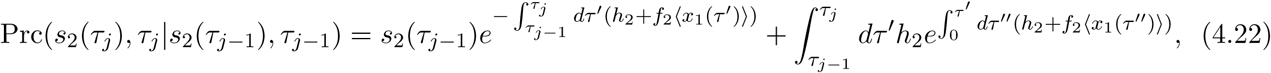

where we have set 〈*Q*_*i*_(*τ*_*j-*1_)〉 = *s*_2_(*τ*_*j-*1_) (with *i* = 0 or 1) expressing the fact that at the instant of time *τ*_*j-*1_ the gene 2 is in state *i* with probability *s*_2_(*τ*_*j-*1_) (which is 0 or 1), as we have done for the gene 1. The difference is that for gene 1 we have closed solutions for the occupancy probabilities and the conditional probabilities were derived analytically from these solutions in (4.17). However, here, as said before, we must perform the integrals in (4.22) numerically.

With (4.21) and (4.22) at hand we are in position to obtain the distributions for mRNA and proteins associated with gene 2 in the same fashion as we did for the gene 1. The only difference is that the mRNA and protein copy numbers will be obtained using (4.15) and (4.19) just by changing the index 1 for 2 (*s*_1_(*τ*) *→ s*_2_(*τ*), *y*_1_(*τ*) *→ y*_2_(*τ*), and *x*_1_(*τ*) *→ x*_2_(*τ*)). A last comment regards the parameter space: as we did for the gene 1, we have redefined the parameter space of gene 2 and introduced the more biological parameters; the asymptotic occupancy probabilities (*q*_0_, *q*_1_) and the flexibility parameter (*σ*). The new parameters are expressed in terms of the old ones as:

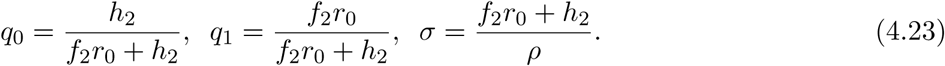

In figure 7 we compare the distributions for mRNA and proteins associated with gene 2 with the direct simulation of the model. We have done that in four distinct situations when the first and the second genes are fast and slow. For all the cases the first and second gene are OFF at *τ* = 0 and both have asymptotic occupancy probabilities equal to 1/2. The comparison is quantitative and uses the distance defined by (4.1).

**Figure 7:**
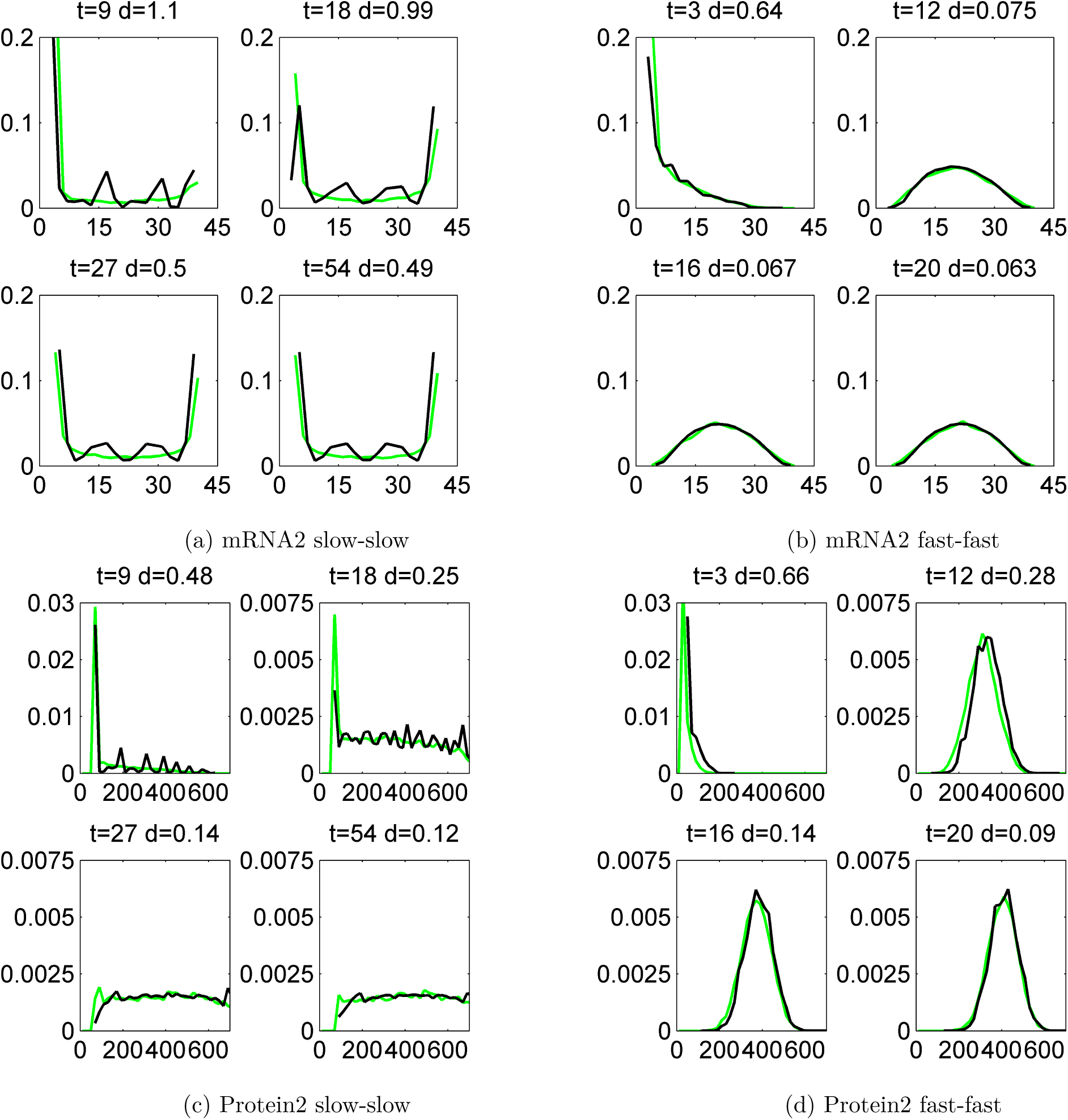
Histograms of protein and mRNA copy numbers for the second gene in the circuit M1, produced by the Monte-Carlo method (green lines) and by the push-forward method (black lines). The initial data is *x*_1_ = *x*_2_ = *y*_1_ = *y*_2_ = 0 and the circuit parameters are *ρ* = 1, *k*_0_ = 4, *k*_1_ = 40, *a* = 1*/*5, *b* = 4, *p*_*a*_ = 0.5, for both genes, and *ε*= 0.5, *ε* = 5.5, for slow, and fast switching genes, respectively. The comparison is quantified by the distance *d* defined by (4.1).

## 5 Testing the propensity of mRNA and protein intrinsic fluctuations to reveal gene/gene interactions

In this paper we used mathematical models to predict mRNA and protein “intrinsic” fluctuations (by “intrinsic” we understand fluctuations generated by the stochastic gene network dynamics). An important question in this context is if “intrinsic” gene expression fluctuations can be used to reverse engineer gene networks. “Extrinsic” fluctuations (by “extrinsic” we understand perturbations of gene networks coming from their environment) of the transcriptome were extensively used in the past to reconstruct gene networks using correlation or mutual information (for a popular method see [31]). A few reverse-engineering studies obtained gene regulation parameters from intrinsic gene expression fluctuations [12, 34]. Quantifying intrinsic fluctuations ask for single cell measurements of several genes. A variety of technologies are now ready to take this challenge: single cell sequencing [46], single-cell RNA microscopy [48], various versions of time-lapse microscopy [33], fluorescence correlation microscopy [12]. It is therefore important to test the propensity of expression fluctuations to discriminate between various gene network architectures.

The predictions of our PDP models are provided as multivariate distributions of mRNAs and proteins copy numbers for one or several genes. These predictions can be directly compared with results obtained with molecular biology and biophysics experimental methods.

First, one would like to test if the differences between distributions predicted with various gene circuit models are significant and therefore can be used to discriminate between gene circuit models. To this end, we generated bivariate proteins and mRNA distributions for five different gene circuits like in Figure 4. The visual inspection of Figure 8, suggests that mRNA distributions resulting from five different gene circuits, with gene/gene interactions that differ by their signs, are very similar in the same regime of switching (fast or slow) of the genes. The mRNA distributions discriminate between model parameters (fast or slow swithcing) but do not discriminate between circuit architectures. The protein bivariate distributions are shown in Figure 9. They differ strongly from mRNA distributions and discriminate between both parameters and architectures.

**Figure 8:**
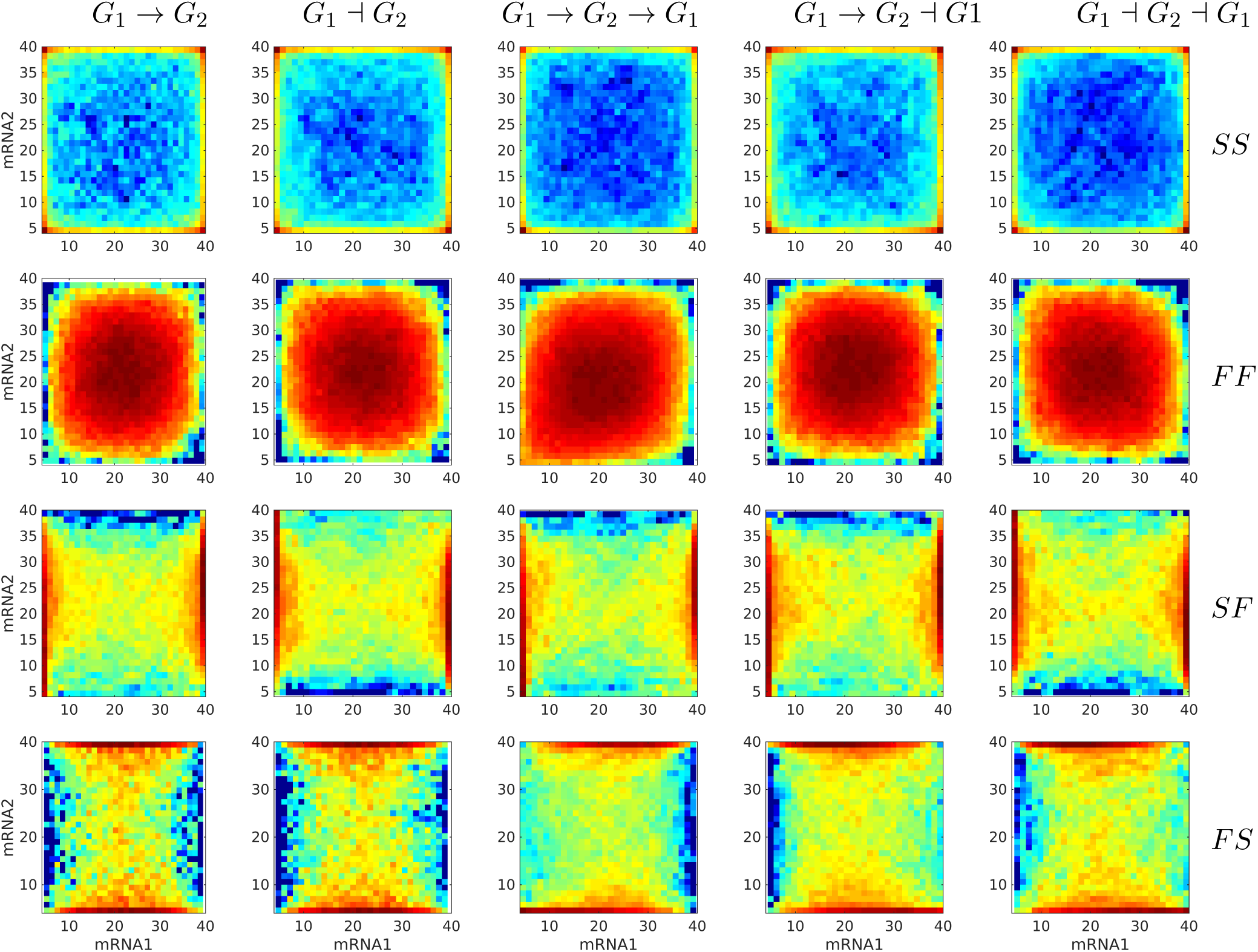
Steady state bivariate histograms of mRNA copy numbers from two interacting genes in circuits of different types and for four switching regimes of the promoters (SS, FF, SF, FS, where SF means that the first gene is slowly switching whereas the second is switching fastly), obtained with the Monte-Carlo method. The individual gene parameters are those used in Figure 5; *f* and *h* constants in *fx*_*i*_ or *hx*_*i*_ terms are chosen such that then mean mRNA and protein are the same in regulated and constitutive genes. The probability to color map relation is logarithmic.

**Figure 9:**
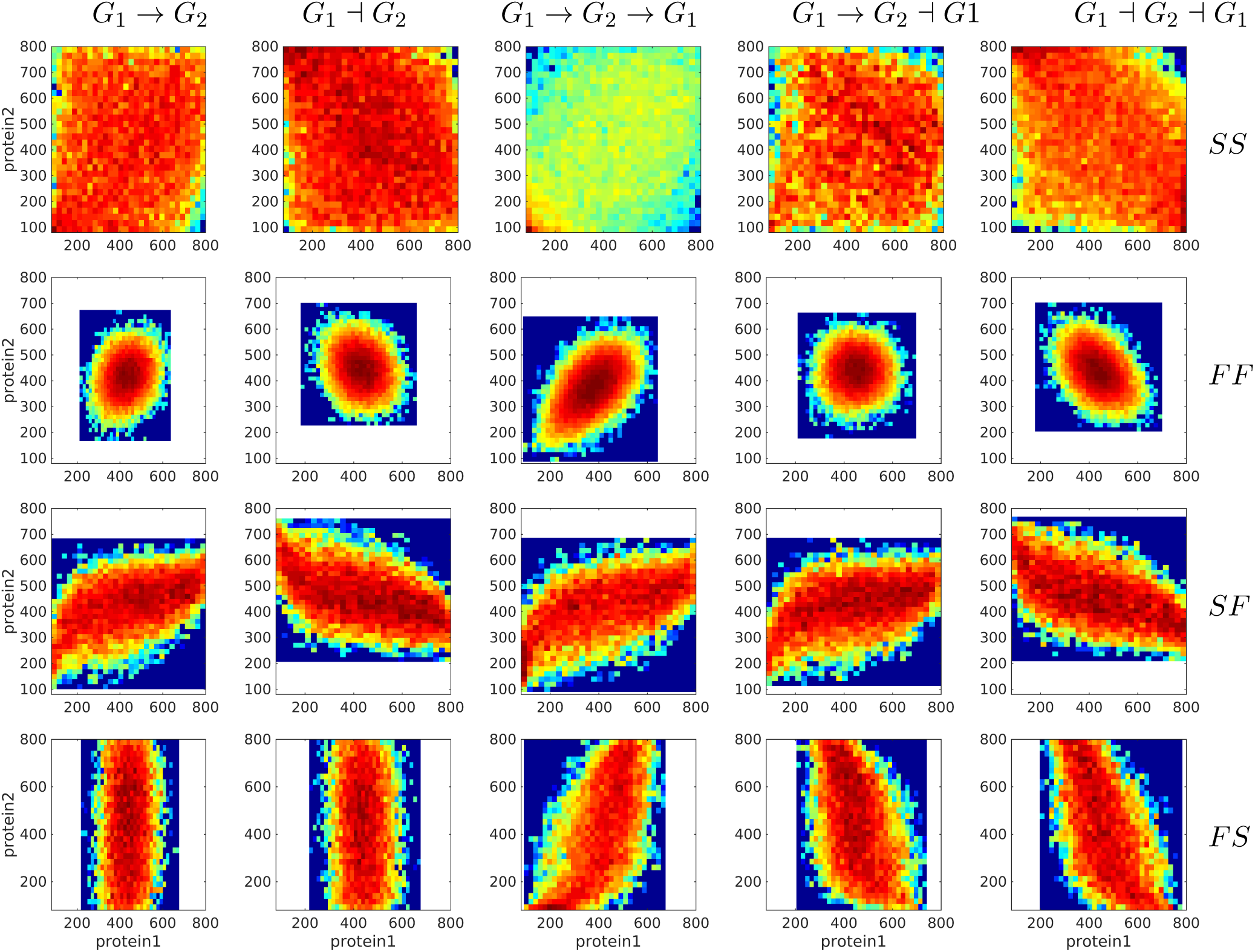
Steady state bivariate histograms of protein copy numbers from two interacting genes in circuits of different types and for four switching regimes of the promoters, obtained with the Monte-Carlo method. The individual gene parameters are those used in Figure 5; *f* and *h* constants in *fx*_*i*_ or *hx*_*i*_ terms are chosen such that then mean mRNA and protein are the same in regulated and constitutive genes. The probability to color map relation is logarithmic.

As visual colormap differences may be judged subjective, we developed a quantitative test for the discriminant power. This test is based on the distance *d* defined by (4.1). We have computed *d* pairwise, for mRNA and for protein distributions produced from different gene circuits at steady state. In order to test if these distances are significantly large we compared them to distribution of distances between random samples generated by the same gene circuit model for a fixed number of cells. The result of this comparison is shown in Figure 10 for the two gene circuits *G*_1_ *→ G*_2_ and *G*_1_ *┤ G*_2_ that differ by the sign of the interaction; one can notice that the protein fluctuation based distance is significant, whereas the mRNA fluctuation distance is not, both for slow/slow and fast/fast genes.

**Figure 10:**
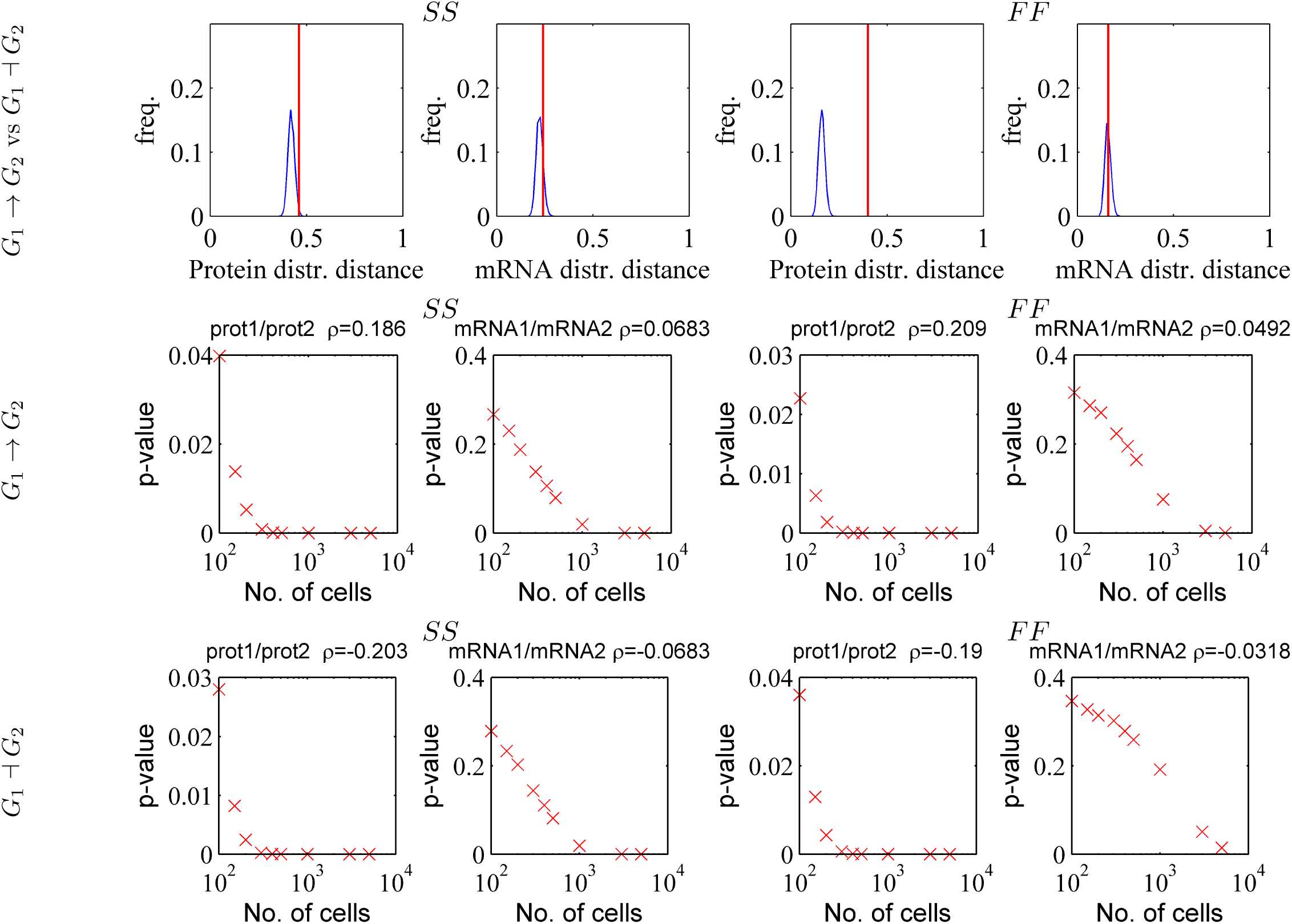
Testing the capacity of mRNA and protein fluctuations to reveal gene/gene interactions. Upper row: the distance between distributions generated by activating and repressive circuit (red line) are compared to the distances between random samples (*N*_*c*_ = 1000) generated by the same repressive circuit (blue smoothed histogram). Middle and lower row: the Bravais-Pearson correlation coefficient *ρ* is computed from random samples containing increasing numbers of cells and the corresponding *p*-value (*p* is the upper tail probability and correlation is significant when *p* is small) is represented vs. sample size. The parameters of the simulations are those used in Figure 9.

We have also tested the significance of the correlation computed from bivariate mRNA or protein distributions. A simple gene reconstruction method is to consider that genes interact if the correlation coefficient is significantly different from zero. We have computed the Bravais-Pearson correlation coefficient from bivariate mRNA and protein samples generated with our PDP model, at steady state and for increasing numbers of cells *N*_*c*_. For both models *G*_1_ *→ G*_2_ and *G*_1_ *┤ G*_2_ a significant (upper tail probability *p <* 5%) protein/protein correlation is obtained for moderate cell populations (*N*_*c*_ *>* 100 for *p <* 5%, see Figure 10). In order to obtain significant mRNA/mRNA correlation one has to use very large numbers of cells (*N*_*c*_ *>* 1000 for *p <* 5%, see Figure 10). This is possible for single cell sequencing and flow cytometry (with the drawback of the lack of precision in estimating the mRNA copy numbers) but is very difficult for techniques such as MS2 tagging microscopy, or time-lapse microscopy.

## Discussion

We have discussed three methods to compute time-dependent distributions of mRNA and protein copy numbers generated by gene networks. All the three methods are much faster than the Gillespie exact chemical master equation. The finite difference Liouville-master equation method is precise and fast for small models. Simple (non-adaptive) finite difference schemes, however, are demanding in terms of space and time resolution leading to computer memory limitations. In future implementations of the Liouville-master equation method we plan to use spectral methods for bypassing these limitations. The push-forward method is not as precise as the Liouville-master equation, but it is much more stable, even at low resolution. The differences between execution times of various methods are illustrated in the Table1 indicating that the push-forward method is the fastest one. The performance of the push-forward method results from the reduced cardinality of the discrete phase space (2 states for one gene, 4 states for 2 genes) which is an improvement with respect to the initial version in [22]. Further improvements, lifting the mean-field approximation for gene coupling will be presented elsewhere.

**Table 1:**
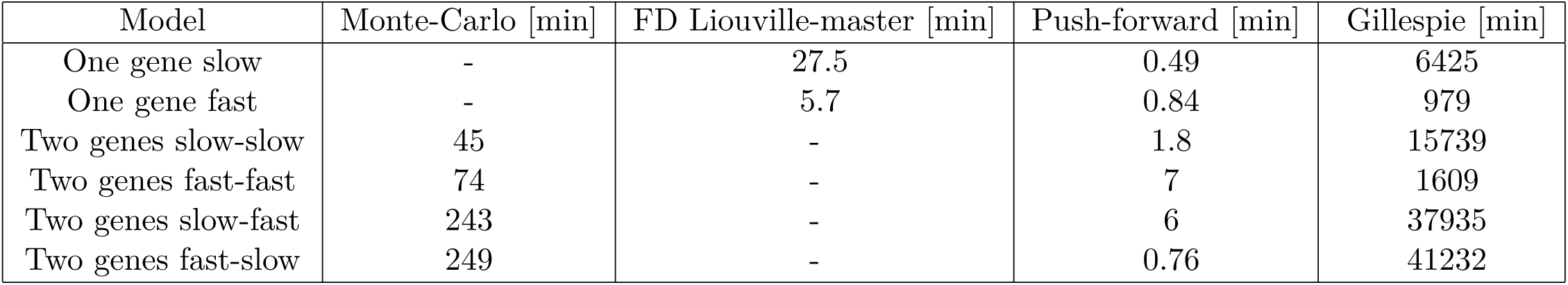
Execution times for different methods. All the methods were implemented in Matlab R2018b running on a single core (multi-threading inactivated) of a Xeon E5 2.4 GHz processor.

| Model | Monte-Carlo [min] | FD Liouville-master [min] | Push-forward [min] | Gillespie [min] |
| --- | --- | --- | --- | --- |
| One gene slow | - | 27.5 | 0.49 | 6425 |
| One gene fast | - | 5.7 | 0.84 | 979 |
| Two genes slow-slow | 45 | - | 1.8 | 15739 |
| Two genes fast-fast | 74 | - | 7 | 1609 |
| Two genes slow-fast | 243 | - | 6 | 37935 |
| Two genes fast-slow | 249 | - | 0.76 | 41232 |

Piecewise-deterministic models are valuable tools for understanding stochastic gene expression in a wide spectrum of regimes, covering both slow and fast switching. The source of stochasticity in such models are the random transitions between discrete promoter states; a phenomenon usually associated with transcriptional bursting. In this paper we have only discussed dichotomous noise (ON/OFF promoters), however, as seen in Section 2, our methods work also for promoters with more than two states.

As application of our numerical methods we tested the capacity of mRNA and protein copy numbers fluctuations to unravel differences between gene circuits architectures. We showed that protein fluctuations are sensitive to differences of architecture and that protein-protein correlation reveals gene-gene interactions for moderate cell population sizes (100 cells). In contrast, mRNA fluctuations are much less sensible to differences in circuit architecture and mRNA-mRNA correlation is small, even for interacting genes. This reinforces the already well established conclusion that proteome contains much more information than the transcriptome. In the past we used the spectrum of protein copy number fluctuations to extract information about promoter repression mechanisms [12]. The difference in behavior of the mRNA and protein fluctuations can be explained by the fact that the mRNA half life is usually much shorter than the protein half life. Gene-gene interactions are mediated by proteins that slowly modulate the gene switching times. Proteins follow these slow modulations, which results in significant protein-protein correlation. mRNA dynamics is permanently submitted to the faster (uncorrelated) gene switching, which explains the lower correlation of steady state mRNA fluctuations. This suggests that reconstruction of gene networks from mRNA intrinsic fluctuations is submitted to severe limitations. More general results concerning these limitations will be presented elsewhere.

## Appendix1: Two gene circuits

The Liouville-master equations for two gene circuits are systems of four PDEs in four variables *x*_1_, *y*_1_, *x*_2_, *y*_2_ representing the protein and mRNA of the first gene *G*_1_ and second gene *G*_2_, respectively. There are five different types of such circuits *G*_1_ *→ G*_2_, *G*_1_ *┤ G*_2_, *G*_1_ *→ G*_2_ *→ G*_1_, *G*_1_ *→ G*_2_ *┤ G*_1_, *G*_1_ *┤ G*_2_ *┤ G*_1_ where *→*, *┤* stand for activation and repression, respectively.

The set of two promoters have four discrete states 1 = (0, 0), 2 = (1, 0), 3 = (0, 1), 4 = (1, 1). The respective probabilities densities are *p*_1_(*x*_1_, *y*_1_, *x*_2_, *y*_2_, *t*), *p*_2_(*x*_1_, *y*_1_, *x*_2_, *y*_2_, *t*), *p*_3_(*x*_1_, *y*_1_, *x*_2_, *y*_2_, *t*), *p*_4_(*x*_1_, *y*_1_, *x*_2_, *y*_2_, *t*).

The corresponding Liouville-master equations are the following: For the circuit *G*_1_ *→ G*_2_.

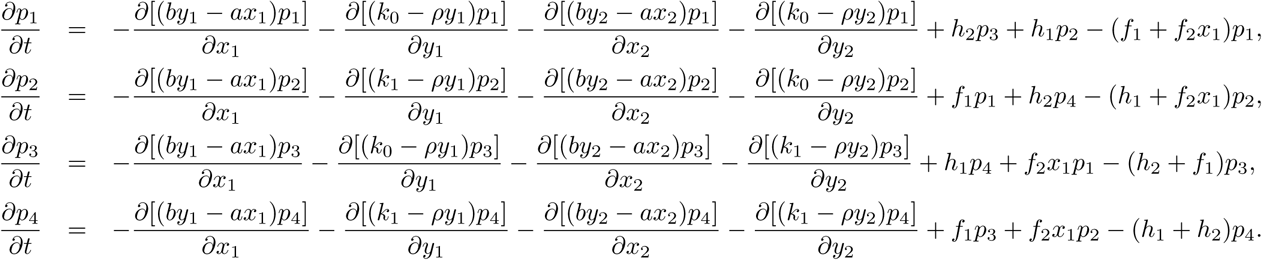

For the circuit *G*_1_ *┤ G*_2_.

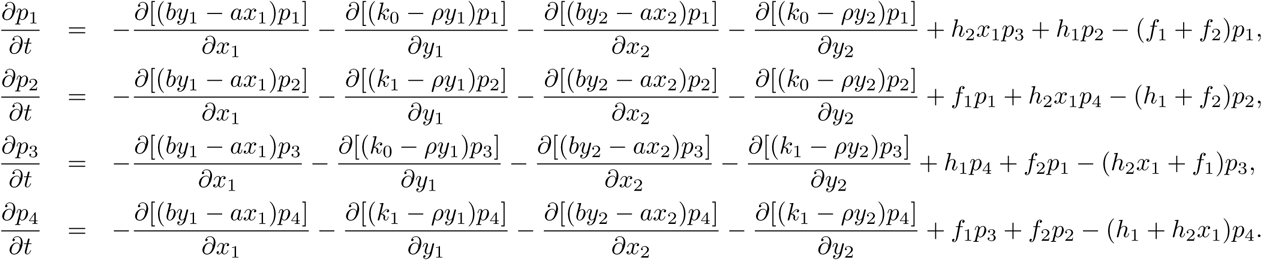

For the circuit *G*_1_ *→ G*_2_ *→ G*_1_.

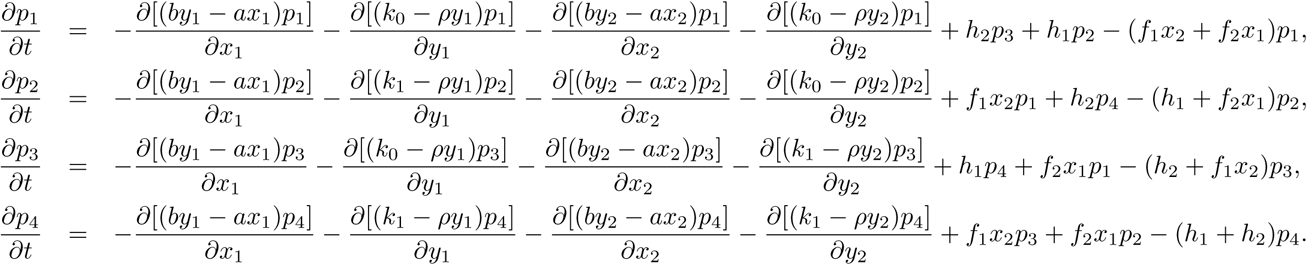

For the circuit *G*_1_ *→ G*_2_ *┤ G*_1_.

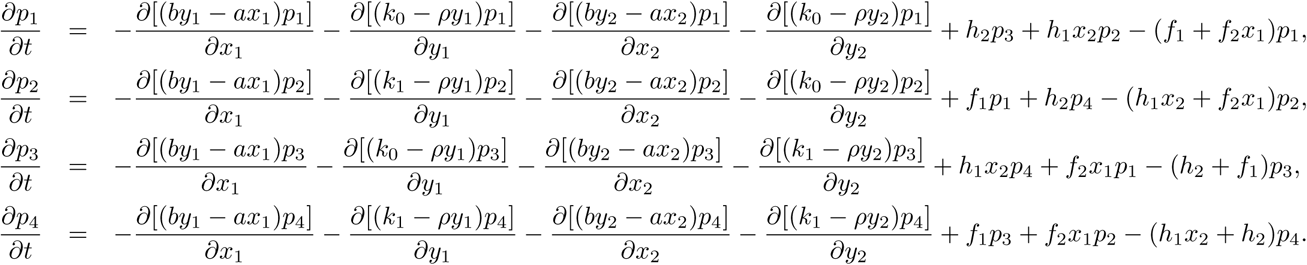

For the circuit *G*_1_ *┤ G*_2_ *┤ G*_1_.

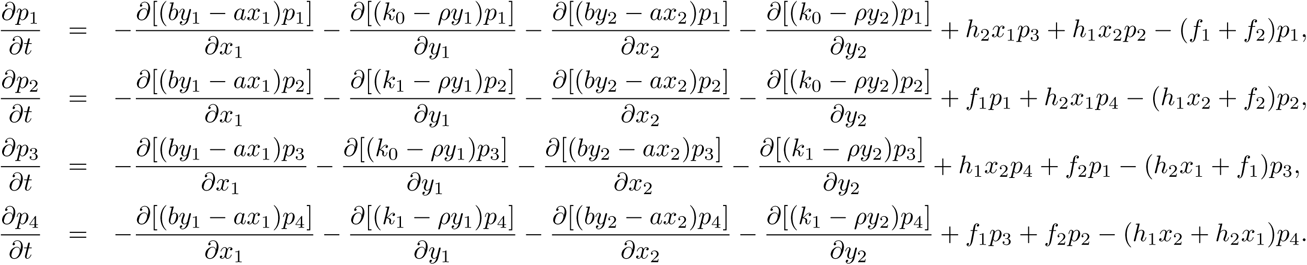

## Appendix2: Analytic solutions for one step deterministic propagation

For gene circuits (4.2) read

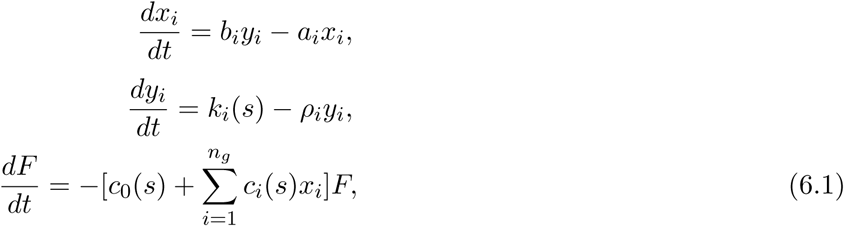

where *c*_*i*_(*s*), *i ∈* [0, *n*_*g*_] are positive constants depending on the discrete Markov chain state *s*.

The solution of this system is straightforward

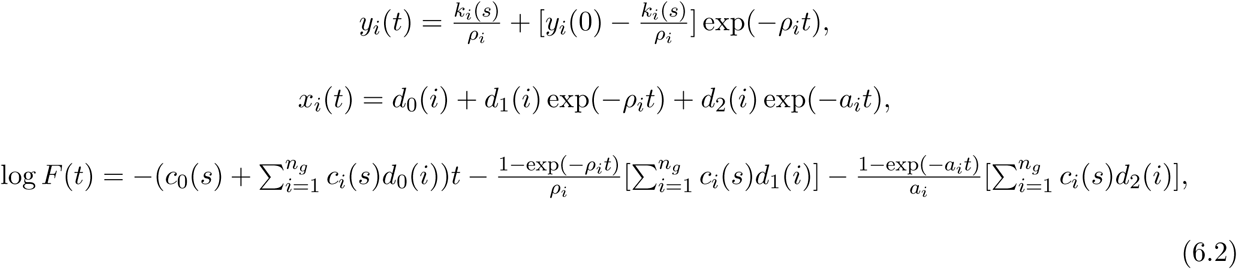

where 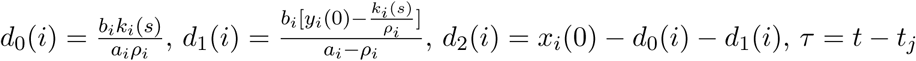

For the single constitutive gene model *c*_0_(0) = *f*, *c*_0_(1) = *h*, *c*_1_(0) = *c*_1_(1) = 0, *F* (*τ*) = exp[−(*f* + *h*)*τ*]. The constants *c*_*i*_, *i ∈ {*0, 1, 2*}* for the two gene models are given in the Table 2. For a general gene network, the constants table is precalculated symbolically and used to generate automatically the simulation code.

**Table 2:** Constants *c*_*i*_ for computing the next step waiting time in the two gene model.

| Model | State | $c_0$ | $c_1$ | $c_2$ |
| --- | --- | --- | --- | --- |
| $G_1 \rightarrow G_2$ | (0,0) | $f_1$ | $f_2$ | 0 |
| | (1,0) | $h_1$ | $f_2$ | 0 |
| | (0,1) | $f_1 + h_2$ | 0 | 0 |
| | (1,1) | $h_1 + h_2$ | 0 | 0 |
| $G_1 \dashv G_2$ | (0,0) | $f_1 + f_2$ | 0 | 0 |
| | (1,0) | $f_2 + h_1$ | 0 | 0 |
| | (0,1) | $f_1$ | $h_2$ | 0 |
| | (1,1) | $h_1$ | $h_2$ | 0 |
| $G_1 \rightarrow G_2 \rightarrow G_1$ | (0,0) | 0 | $f_2$ | $f_1$ |
| | (1,0) | $h_1$ | $f_2$ | 0 |
| | (0,1) | $h_2$ | 0 | $f_1$ |
| | (1,1) | $h_1 + h_2$ | 0 | 0 |
| $G_1 \rightarrow G_2 \dashv G_1$ | (0,0) | $f_1$ | $f_2$ | 0 |
| | (1,0) | 0 | $f_2$ | $h_1$ |
| | (0,1) | $f_1 + h_2$ | 0 | 0 |
| | (1,1) | $h_2$ | 0 | $h_1$ |
| $G_1 \dashv G_2 \dashv G_1$ | (0,0) | $f_1 + f_2$ | 0 | 0 |
| | (1,0) | $f_2$ | 0 | $h_1$ |
| | (0,1) | $f_1$ | $h_2$ | 0 |
| | (1,1) | 0 | $h_2$ | $h_1$ |

## Appendix3: Supplementary information for the push-forward method

In order to obtain an expression for 〈*x*_1_(*τ*)〉 we must solve a set of ordinary differential equations:

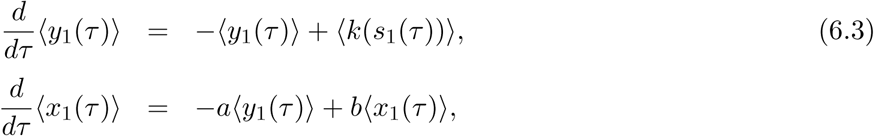

where 〈*k*(*s*_1_(*τ*))〉 = *k*_0_*P*_0_(*τ*) + *k*_1_*P*_1_(*τ*). The solution for 〈*y*_1_(*τ*)〉 is:

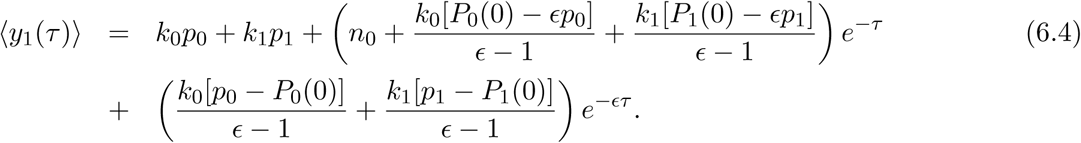

The general solution for protein mean value has the structure, 〈*x*_1_(*τ*)〉 = *r*_0_ + *r*_1_*e*^*-τ*^ + *r*_2_*e*^*-aτ*^ + *r*_3_*e*^*-Eτ*^, where the coefficients are given by:

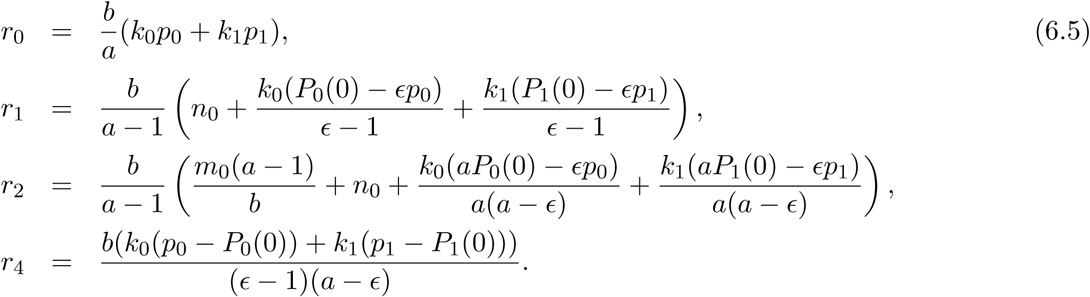

